# petVAE: A Data-Driven Model for Identifying Amyloid PET Subgroups Across the Alzheimer’s Disease Continuum

**DOI:** 10.64898/2026.02.02.703218

**Authors:** Arina A. Tagmazian, Claudia Schwarz, Catharina Lange, Esa Pitkänen, Eero Vuoksimaa, the Alzheimer’s Disease Neuroimaging Initiative

## Abstract

Amyloid-β (Aβ) PET imaging is a core biomarker and is sufficient for the biological diagnosis of Alzheimer’s disease (AD). Here, we aimed to identify biologically meaningful subgroups across the continuum of Aβ accumulation using a data-driven deep learning approach, without imposing predefined thresholds for Aβ negativity or positivity.

We analyzed 3,110 Aβ PET scans from the Alzheimer’s Disease Neuroimaging Initiative and Anti-Amyloid Treatment in Asymptomatic Alzheimer’s Disease studies to develop petVAE, a two-dimensional variational autoencoder. The model accurately reconstructed scans without prior labeling, selection by scanner or region of interest. Latent representations of scans extracted from petVAE were used to visualize and cluster the AD continuum. Clustering yielded four groups: two predominantly Aβ negative (Aβ-, Aβ-+) and two predominantly Aβ positive (Aβ+, Aβ++).

All clusters differed in standardized uptake value ratio (*p* < 1.64×10⁻⁸) and cerebrospinal fluid (CSF) Aβ (*p* < 0.02), demonstrating petVAE’s ability to assign scans along the Aβ continuum. Extreme clusters (Aβ-, Aβ++) resembled conventional Aβ negative and positive groups and differed in cognition, *APOE* ε4 prevalence, Aβ and tau CSF biomarkers (*p* < 3×10⁻⁶). Intermediate clusters (Aβ-+, Aβ+) showed higher odds of carrying at least one *APOE* ε4 allele versus Aβ- (*p* < 0.03). Participants in Aβ+ or Aβ++ clusters exhibited faster progression to AD (Aβ+ hazard ratio = 2.42, Aβ++ HR = 9.43; *p* < 1.17×10⁻⁷).

Thus, *petVAE* was capable of reconstructing PET scans while extracting latent features that capture the AD continuum and define biologically meaningful subgroups, enabling data-driven characterization of preclinical disease stages.

## 1. Introduction

The early diagnosis of Alzheimer’s disease (AD) and prediction of its development remain among the most relevant questions in neuroscience. Even in early stages, annual costs per patient exceed €16,000 and rise with AD progression [1]. As AD accounts for 60–80% of dementia cases, it poses a major socio-economic issue [2]. Although subtle cognitive changes, including episodic memory impairment, may occur early in the disease course and can in some cases predict underlying pathology [3,4], these measures remain neither sufficiently sensitive nor specific for AD [5,6]. In contrast, AD biomarkers represent an earlier and more disease-specific process, making it a key target for improving diagnostic accuracy.

In the AT(N) research framework, where A - amyloid-β (Aβ) aggregation, T - tau pathology, and N - neurodegeneration [7,8], Aβ biomarker is the earliest to appear in the brain and is considered a Core 1 biomarker that has been suggested to be sufficient for the biological diagnosis of AD and to inform clinical decisions [9]. Owing to its early and disease-specific accumulation, Aβ assessment with positron emission tomography (PET) provides valuable information on both the magnitude and spatial distribution of amyloid burden in the brain [8]. Conventional PET quantification approaches, such as the standardized uptake value ratio (SUVR) and centiloid (CL) scaling, summarize global cortical Aβ burden and place individuals along a continuous AD spectrum, which is typically stratified into discrete groups using predefined thresholds. Due to the limited comparability of SUVR across tracers and datasets, the CL scale was developed to enable standardization [10]. However, despite its advantages, the CL framework remains inherently threshold-dependent, and these thresholds can vary across studies, particularly in the early and intermediate stages of Aβ accumulation [11]. This limitation highlights a potential methodological advantage of the data-driven deep learning (DL) approaches, which can model and group the Alzheimer’s disease continuum without relying on predefined thresholds [12].

DL has been widely applied in AD research for detection, stage prediction, and neuroimaging tasks such as segmentation and spatial normalization [13,14]. More recently, unsupervised approaches, including variational autoencoders (VAEs) [15], have been explored for brain anomaly detection, disease severity indexing, metabolomic change prediction and tau pathology progression [16–18]. Several DL studies have identified AD subgroups from Aβ and tau PET data using both supervised and unsupervised methods and compared them across demographic and cognitive measures [19–21]. However, most of these approaches aimed to define clusters that are distinctly separated from one another, whereas AD pathology develops along a continuum. Studies using SUVR and CL scaling have shown that continuum-based representations of AD with finer stratification improve the prediction of cognitive decline and disease progression compared with more discrete, binary Aβ classification [22,23].

In this work, we apply VAEs to Aβ PET data to learn latent representations that capture this continuous spectrum of amyloid aggregation, enabling the discovery of data-driven subgroups without relying on predefined diagnostic boundaries. We used a 2D VAE to define data-driven AD subgroups from Aβ PET scans. We then examined their associations with cerebrospinal fluid (CSF) biomarkers, cognitive performance, genetic risk, and progression to AD. We also evaluated their agreement with SUVR- and CL-based grouping. Such an approach could streamline clinical practice by automatically assigning patients to disease subgroups and linking them to neuropsychological measures, Apolipoprotein E (*APOE*) genotype (ε4-carrier versus non-carrier), and to biomarkers. Additionally, DL-based representations provide an alternative, data-driven characterization of tracer uptake patterns that may complement established SUVR- and CL-based measures.

## 2. Methods and materials

### 2.1. Data

In total, we used 3,110 [^18^F]-AV45 (Florbetapir) Aβ PET scans from Alzheimer’s Disease Neuroimaging Initiative (ADNI) and Anti-Amyloid Treatment in Asymptomatic Alzheimer’s Disease (A4/LEARN) studies for training and testing the model (Suppl. Table 1). To test model generalizability across tracers, we used 344 [^18^F]-FBB (Florbetaben) and 119 [^11^C]-PiB (Pittsburgh compound B) Aβ PET from ADNI.

#### 2.1.1. ADNI

Data used in the preparation of this article were obtained from the Alzheimer’s Disease Neuroimaging Initiative (ADNI) database (adni.loni.usc.edu). The ADNI was launched in 2003 as a public-private partnership, led by Principal Investigator Michael W. Weiner, MD. The primary goal of ADNI has been to test whether serial magnetic resonance imaging (MRI), PET, other biological markers, and clinical and neuropsychological assessment can be combined to measure the progression of mild cognitive impairment (MCI) and early Alzheimer’s disease (AD). [24] The current goals include validating biomarkers for clinical trials, improving the generalizability of ADNI data by increasing diversity in the participant cohort, and to provide data concerning the diagnosis and progression of Alzheimer’s disease to the scientific community. For up-to-date information, see adni.loni.usc.edu.

We used 1,782 Aβ PET scans from the ADNI dataset, along with corresponding T1 MRI scans taken either at the same visit or within one year of the PET scan. The scans were not pre-selected by AD stage, ADNI phase or scanners. This data was used for model training, validation, and testing and included scans obtained only with AV45 tracer. Among them there were 495 scans from cognitively normal patients, 163 from subjective memory concerned patients, 967 from those with mild cognitive impairment and 157 from patients with AD (Suppl. Table 1). A total of 344 FBB and 119 PiB PET scans were used as external validation data to assess the model’s ability to generalize across different amyloid tracers. For further co-registration and model training we used scans preprocessed by ADNI (‘Co-registered, averaged’), where six five-minute frames were averaged in native space.

#### 2.1.2. A4/LEARN

We used 1,328 AV45 Aβ PET scans from the A4/LEARN dataset and corresponded T1 MRI scans. All PET scans were reconstructed into 4×5-minute frames.

### 2.2. Methods

#### 2.2.1. Data pre-processing

In our downstream analysis we used composite SUVR, CSF biomarker measurements (Aβ42, tau and pTau181) and the Preclinical Alzheimer Cognitive Composite (PACC) score. For ADNI data, SUVR was calculated as the ratio of the cortical summary region (frontal, anterior/posterior cingulate, lateral parietal, and lateral temporal) to the whole cerebellum, with an Aβ-positive threshold of 1.11 [25,26]. For A4 scans, the cortical summary region included the frontal, temporal, parietal cortices, precuneus, and anterior and posterior cingulate, with the whole cerebellum as a reference region and a threshold of 1.10 for Aβ-positivity [27,28].

PACC scores were calculated using the Free and Cued Selective Reminding Test (FCSRT), Logical Memory II Story A Delayed Recall (LM), Digit Symbol Substitution Test (DSST), and Mini-Mental State Examination (MMSE) for the A4 dataset. For the ADNI dataset, the FCSRT was replaced with Delayed Word Recall from the Alzheimer’s Disease Assessment Scale–Cognitive Subscale [29].

CSF measures were available only for the ADNI dataset and exhibited right-skewed distributions; therefore, we used natural log-transformed values standardized to a mean of 0 and a standard deviation of 1 instead of the original values. Also, some Aβ CSF measurements had a ‘>1,700’ label, and we excluded them from analysis due to missing original values (Suppl. Fig. 1). SUVR values were also log-transformed and standardized in the same way, whereas PACC scores were only z-transformed.

#### 2.2.2. Aβ PET scans pre-processing

All Aβ PET scans were co-registered first to a corresponding individual T1 MRI image and after that to the T1 MNI152 template using the affine transformation option in the ANTsPy python package [30].

In addition, A4/LEARN scans required frame-to-frame registration that was implemented with ANTsPy (‘Rigid’ transformation) before common PET-MRI-MNI152 co-registration. We averaged all frames and created an individual PET template, co-registered each frame to the template and averaged the already registered frame. These averaged PET scans were used for the MRI registration.

For quality control of PET-to-MRI registration, we used Pearson correlation coefficient between voxel intensities of the two images from the ‘image_similarity’ function of ANTsPy package. Based on visual inspection of the data, we set a Pearson correlation threshold of 0.6 and included only scans meeting or exceeding this value.

It should be noted that, in the current study, the term “reconstructed PET scan” refers to the output of the petVAE model, which deconstructs and reconstructs the original PET images, and does not relate to the reconstruction of raw PET data.

All Aβ PET scans were normalised to the reference region (whole cerebellum). We used Hammers brain atlas to define reference regions on scans [31,32] and adjusted standardized uptake value (SUV) values to the reference (SUVR). To mitigate the long-tailed distribution of Aβ PET SUVR values, we applied clipping at the 0.999 quantile. The resulting values were subsequently rescaled to the [0,1] interval to standardize the range across samples.

#### 2.2.3. petVAE model’s architecture and parameters

The petVAE is a 2D convolutional variational autoencoder (VAE) [15] that contains 1.10 million parameters (Fig. 1). A VAE learns to compress an input image (here, a PET scan slice) into a small set of latent variables that capture its most important underlying features needed to reconstruct the input image. It is trained by reconstructing the original image from this compressed representation, minimizing both the reconstruction error and a regularization term that encourages the latent variables to follow a predefined prior distribution, such as Gaussian.

**Figure 1.**
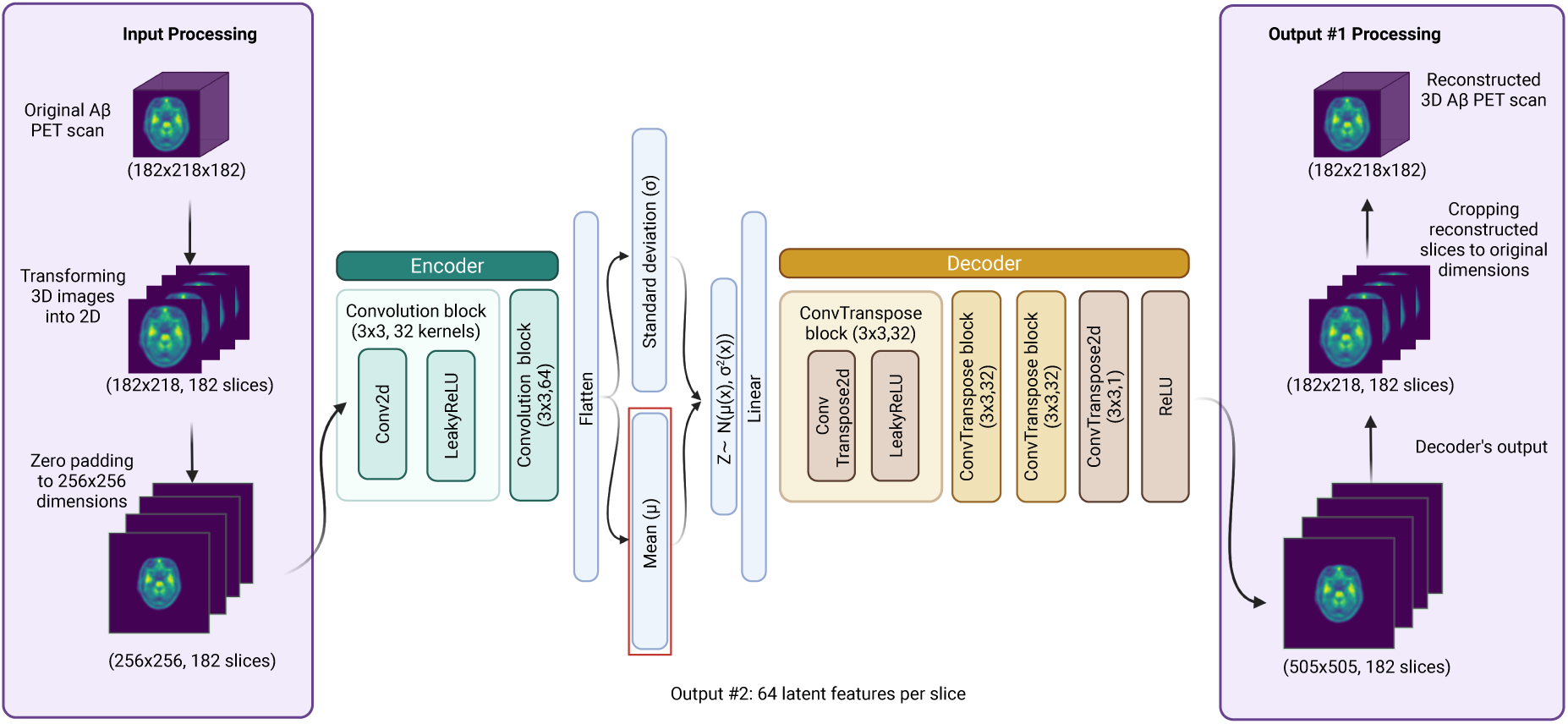
Schematic representation of the petVAE model architecture. “(3×3, 32 kernels)” indicates the kernel size and number of kernels used in the convolutional (Conv) and deconvolutional (ConvTranspose) layers. ReLU and LeakyReLU were used as activation functions [33,34]. Created with BioRender.com.

The Aβ PET scans, each with a size of 182×218×182 voxels, were split into 182 2D axial slices and used as petVAE input. Each slice was masked using the Hammers brain atlas to exclude the skull and background [31]. To standardize input dimensions, slices were centered and zero-padded to 256×256 to make the model applicable across scans of varying original sizes. Eighty percent of the pooled and mixed ADNI and A4 datasets (n = 2,489) was allocated to model development. From these, 80% were used for training (n = 1,991) and 20% for validation (n = 498). The remaining 20% of the dataset was reserved for testing the final model (n = 621). The model was trained on approximately 360,000 Aβ PET slices.

The petVAE outputs both the reconstructed Aβ PET slices and 64 latent features per slice. The latent features are then aggregated across all slices to form a single 11,648-dimensional feature vector for each scan. We used the Adam optimization algorithm (Kingma and Ba 2014) with a learning rate of 0.00001 to iteratively update the model parameters by minimizing the loss function during training. The learning rate controls the step size of each update to ensure stable and gradual convergence. The model was trained for up to 300 epochs with a minibatch size of 64 Aβ PET slices, with early stopping applied if the loss evaluated in the validation dataset did not improve for 10 consecutive epochs. One epoch corresponds to a single pass through the entire training dataset. The loss function (Suppl. Eq. 1) quantifies the discrepancy between the reconstruction and the original scan with mean squared error (MSE) (Suppl. Eq. 2), and the deviation of the latent variables from the prior with Kullback-Leibler divergence (KLD) (Suppl. Eq. 3) [15]. Additionally, reconstruction accuracy was also evaluated by the structural similarity index measure (SSIM) (Suppl. Eq. 4) and peak signal-to-noise ratio (PSNR) (Suppl. Eq. 5). The petVAE code is available at GitHub (https://github.com/artagmaz/petVAE).

#### 2.2.4. Selecting petVAE latent features defining the AD continuum based on association with AD biomarkers

In all subsequent analyses, we excluded scans from participants with clinical diagnosis of AD, allowing us to focus on earlier stages of amyloid aggregation. Because our main objective was to study amyloid-related features, we classified scans into SUVR-defined Aβ- and Aβ+ groups, and used a Random Forest (RF) model on the petVAE-derived latent features to classify scans either as Aβ- or Aβ+. To run the RF model and all subsequent feature-based analyses we used all available scans from the training, validation, and test subsets. We then extracted impurity-based feature importances (mean decrease in impurity, MDI) from the RF model, where “impurity” reflects how mixed the class labels (Aβ-vs. Aβ+) are within the data, and features that more strongly reduce this mixing when used for splitting are considered more important. Thus, we were able to quantify the contribution of each latent feature to amyloid status differentiation.

First, we identified features that accounted for 95% of total importance (‘Core 95% features’), as well as the features with an importance score greater than 0.01 (‘high-impact features’). To visualize high-impact features and brain areas that they represent, we computed Pearson correlation maps. More detailed methodology is described in the Supplementary file. Due to the large number of features in the core 95% feature set (n = 3,098), voxel-wise correlation mapping was not performed for it.

We used Uniform Manifold Approximation and Projection (UMAP), a nonlinear dimensionality reduction technique, to generate low-dimensional embeddings from the selected features and to rank each scan along the AD continuum based on its position in the embedding space (Suppl. Fig. 2). Ranking was performed by fitting a parabola to the UMAP coordinates, projecting each point onto the curve, and ordering them along its arc length. Specifically, for each scan, candidate projection points were sampled along the parabola near its x-coordinate, and the closest point (based on Euclidean distance) was selected as the projection. The projected points were then ordered along the curve by increasing x-coordinate, such that the projection with the minimum x-value defined the starting point of the trajectory. Ranks were assigned based on cumulative distance along the parabola from this point. Although referred to as “ranking,” this metric is continuous rather than ordinal: the cumulative distances were normalized to the [0,1] interval, where 0 and 1 correspond to the minimum and maximum positions along the trajectory in the dataset, respectively. Thus, higher values reflect progressively later positions along the derived trajectory (Suppl. Fig. 2).

The UMAP structure and rank stability were assessed using bootstrap resampling (n = 1,000). For each iteration, a random subset of 50% of samples was used to compute UMAP embeddings and trajectory ranks. For each sample, the mean rank and standard deviation across bootstrap runs were then calculated to quantify positional stability (Suppl. Fig. 3).

To investigate the association between scan ranks and relevant clinical and imaging variables, we calculated Spearman correlations between scan ranks and CSF biomarkers, SUVR values, and PACC score. In addition, we fitted linear regression models with scan ranking as the independent variable and age and sex as covariates to predict CSF biomarker concentrations, SUVR values, and PACC score. For *APOE* ε4 genotype, we conducted two separate logistic regression analyses with identical covariates. One model contrasted *APOE* ε4 heterozygous individuals with non-carriers, and the second model contrasted *APOE* ε4 homozygous individuals with non-carriers.

#### 2.2.5. Clustering of AD continuum and exploring clusters’ association with AD biomarkers

To define clusters of scans that are the most similar by selected features we applied hierarchical clustering to divide the scans into two, three, and four clusters (Suppl. Table 2, Suppl. Fig. 8). We compared the results of petVAE-based clustering with SUVR-based groups using a confusion matrix. However, rather than treating the SUVR groups as ground truth and computing accuracy, we focused on assessing the agreement between the two grouping methods. Therefore, we report metrics such as concordance (Suppl. Eq. 1) and positive/negative agreement (Suppl. Eq. 2,3). In the two-cluster solution, the cluster with lower amyloid burden was labeled Aβ- and the cluster with higher burden Aβ+. In the three- and four-cluster solutions, intermediate clusters were retained as separate intermediate clusters (Aβ-+ and Aβ+), while Aβ- and Aβ++ were assigned to the clusters with the lowest and highest amyloid burden, respectively.

To compare clusters with each other, we built linear regression models with cluster identifier, age, and sex as covariates and CSF biomarker concentrations, SUVR values, and PACC scores as dependent variables. For *APOE* ε4 risk alleles we used logistic regression with the same covariates. In addition to testing whether the mean levels of the AD-related measures differed between clusters, the models allowed us to estimate effect sizes with confidence intervals, assess the statistical significance of group differences after adjusting for age and sex, and evaluate the relative contribution of each covariate to the variance explained. Only baseline samples were used in the regression models, while subsequent survival analyses were performed on the same baseline cohort using longitudinal follow-up data available in ADNI.

To contextualize our model-based clustering within established approaches to the AD continuum, we compared our results with CL-based stratification. Specifically, we adopted previously defined thresholds [23] to available CL values from ADNI dataset, assigning scans with CL < 15 to the Aβ-reference group, 15–25 CL to Aβ-+, 26–50 CL to Aβ+, and >51 CL to Aβ++. We then repeated the regression and survival analyses using these CL-based groups and compared how well each clustering approach captured differences in biomarker levels, cognitive performance, and disease progression. To quantitatively evaluate and compare clustering strategies, we additionally performed a 10-fold cross-validation framework to assess out-of-sample predictive performance of Centiloid, Core 95%, and high-impact features-based clustering approaches. For continuous outcomes (CSF biomarkers, SUVR values, and PACC scores), predictive performance was quantified using the coefficient of determination (R²), whereas *APOE* ε4 status was evaluated using classification accuracy. In each fold, models were trained on 90% of the data and tested on the remaining 10%, and performance metrics were aggregated across folds. Group-level differences between clustering strategies were assessed using paired t-tests, and multiple comparisons were controlled using false discovery rate (FDR) correction via the Benjamini-Hochberg procedure.

#### 2.2.6. Survival analysis

We used a survival analysis to evaluate AD progression and compare it between clusters. Baseline cognitively healthy and mild cognitive impairment participants from ADNI (n = 574) with available follow-up of up to 72 months were included. Time-to-event was defined as the interval from baseline to diagnosis of AD, and individuals without progression during follow-up were right-censored at their 72 months follow-up visit. We estimated survival curves using the Kaplan-Meier method and compared them between clusters using the log-rank test. In addition, Cox proportional hazards models were fitted with cluster membership as the main predictor and age as well as sex as covariates to account for potential confounding. Hazard ratios with 95% confidence intervals were reported.

#### 2.2.7. Programming environment

Python 3.7.12 and PyTorch 1.13.1 installed with CUDA 11.7 support were used to develop the machine learning models and perform computational analyses [35,36]. The models were trained on NVIDIA Tesla V100 GPUs with 16 GB memory. For processing PET images in Python, we used the Nilearn, nibabel, DLTK, and ANTsPy packages [30,37–39]. PET scan files were converted from DCM to NIfTI format using the dcm2niix package when necessary [40]. All scripts used in the study are provided at https://github.com/artagmaz/petVAE.

During the preparation of this manuscript, the authors used ChatGPT (OpenAI) to assist with language editing and improving the clarity of the text. The authors take full responsibility for the content of the manuscript.

## 3. Results

### 3.1. Performance of the petVAE model

We developed and successfully trained petVAE model to reconstruct axial slices of Aβ PET scans obtained from different scanners with high accuracy (Methods 2.2.3.). The model training converged after 257 epochs. For the test dataset, the model reconstructed brain regions with an average SSIM of 0.90 and a PSNR of 29.6 dB, indicating high structural and intensity fidelity (Fig. 2). SSIM ranges from 0 to 1, with higher values indicating greater structural similarity to the reference image. PSNR is expressed in decibels (dB), with higher values corresponding to better reconstruction quality; values around 20–30 dB are typically considered moderate, while values above 30 dB indicate high-quality reconstruction. The model was used to extract a total of 11,648 latent features for each of the 3,110 scans.

**Figure 2.**
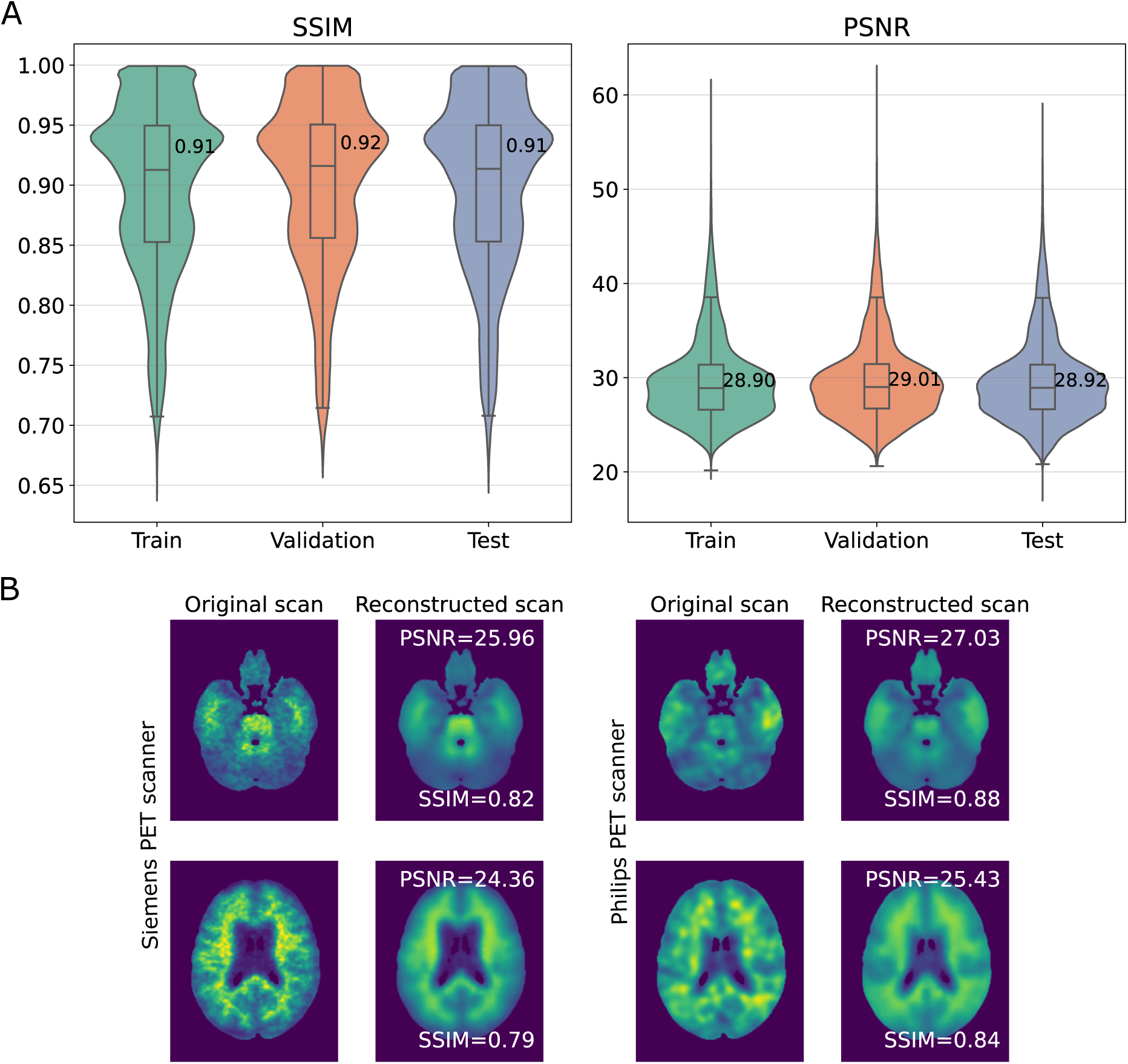
Evaluation of model performance. (A) Distribution of structural similarity index measure (SSIM) and peak signal-to-noise ratio (PSNR) values for the train, validation and test datasets. (B) Examples of PET slices from test dataset and their reconstructions by petVAE. PET scan on the left was acquired with a Siemens scanner, on the right with a Philips Medical Systems scanner. The Philips and Siemens scans are shown as illustrative examples to demonstrate how petVAE handles differences in image resolution across scanners.

### 3.2. Optimal features set for accurate definition of AD continuum

We assumed that not all of the 11,648 latent features extracted per scan are associated with amyloid aggregation. To focus on the most informative variables for subsequent analyses, we defined three feature subsets: all features (n = 11,648), features that cover 95% of Random Forest total importance (‘Core 95% features’, n = 3,098) and the top 10 features according to the importance score (‘high-impact features’, n = 10, importance score > 0.01) (Fig. 3A).

**Figure 3.**
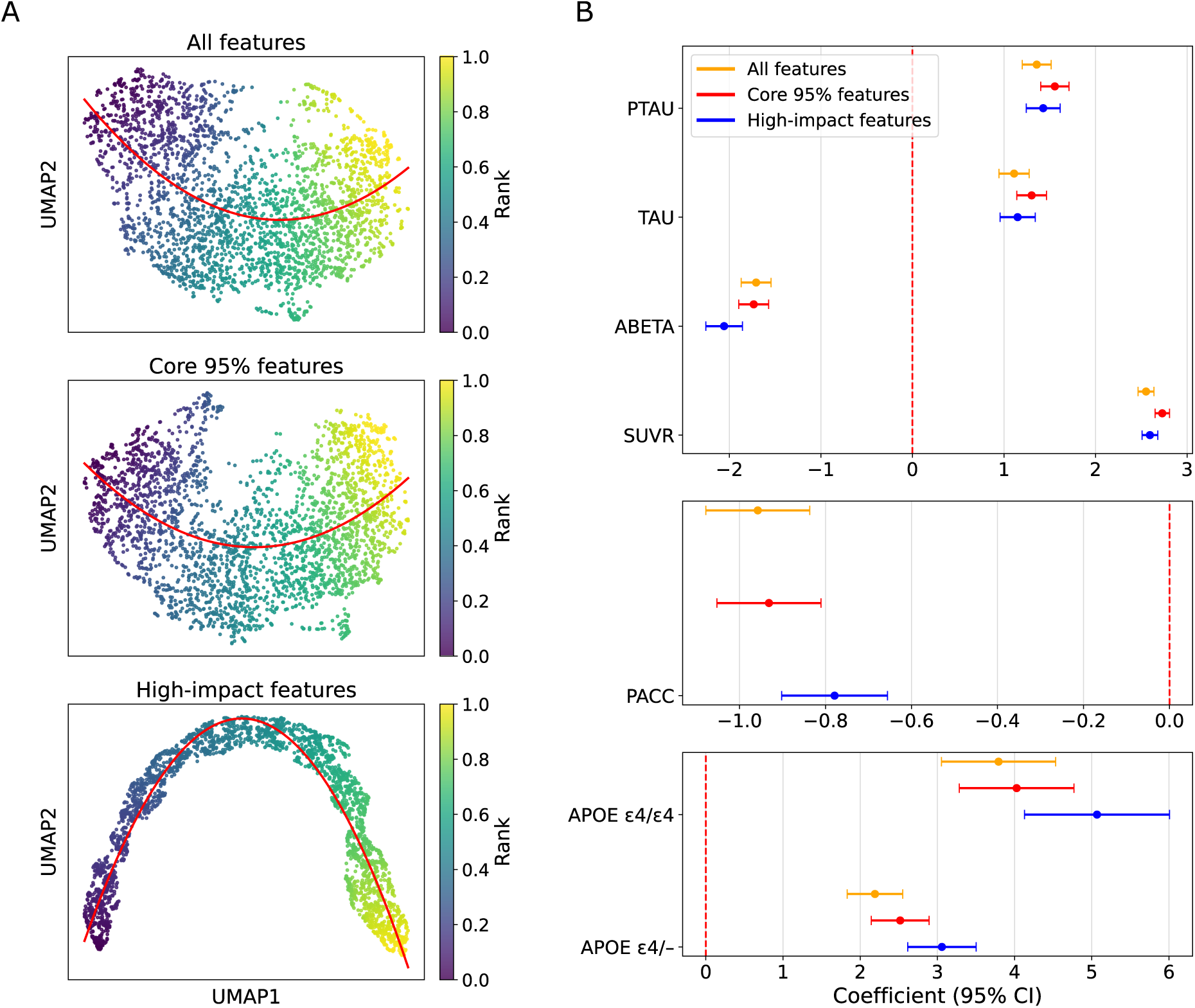
(A) UMAPs embeddings for samples based on all features, core 95% features and high-impact features colored by assigned rank. Red curve – fitted to the UMAP parabola. (B) Associations between ranking variables and AD-related biomarkers. Abbreviations: ABETA – Aβ cerebrospinal fluid biomarker, TAU – tau cerebrospinal fluid biomarker, PTAU – phosphorylated tau cerebrospinal fluid biomarker, SUVR – composite standardized uptake value ratio, *APOE* ε4/– - *APOE* ε4 heterozygous, *APOE* ε4/ε4 - *APOE* ε4 homozygous, PACC - Preclinical Alzheimer’s Cognitive Composite.

We constructed UMAP embeddings for all three feature subsets, fitted a parabolic function to the resulting manifolds, and assigned ranks to the samples based on their proximity to the fitted parabola (Suppl. Fig. 2). The stability of rank values was examined by UMAP and bootstrapping. For both the Core 95% and high-impact feature sets, rank variability increased toward the extremes of the continuum, whereas ranks around 0.5 were the most stable. The high-impact feature set showed a more concentrated and structured distribution of rank deviations, indicating a more consistent representation of the continuum. In contrast, the core 95% feature set exhibited greater dispersion. The model using all features showed the highest variability overall, with a less defined structure, suggesting reduced stability compared to the more selective feature sets (Suppl. Fig. 3).

The ranking of scans was strongly associated with SUVR, CSF biomarkers and PACC scores, adjusting for age and sex (*p* < 0.0005) (Fig. 3B, Suppl. Fig. 4). All three feature subsets demonstrated strong associations between ranks and SUVR, with the core 95% features set yielding the highest coefficient of determination (R² = 0.629, *p* < 1×10^-100^), followed by the high-impact features set (R² = 0.566, *p* < 1×10^-100^) and the all features set (R² = 0.552, *p* < 1×10^-100^). Associations with CSF biomarkers were moderate. The core 95% features set had the highest R² values for Aβ, tau and pTau (R²_Aβ_ = 0.368, *p* = 1.11×10⁻⁷⁶; R²_tau_ = 0.237, *p* = 1.21×10^-49^; and R²_pTau_ = 0.313, *p* = 3.51×10^-73^). Although the core 95% features set achieved the best overall fit, the high-impact features set showed the strongest effect of ranking on Aβ prediction (β = −2.057, *p* = 4.71×10^-73^). Associations with PACC scores were weak across all three feature subsets (0.105 < R² < 0.131, *p* < 1.76×10^-34^).

After adjusting for age and sex, logistic regression revealed that higher ranking was associated with an increased probability of carrying one or two *APOE* ε4 risk alleles (Fig. 3B). The strongest effect of ranking was observed in the high-impact features set (β = 3.058 for *APOE* ε4 heterozygous, *p* = 9.95×10^-42^; β = 5.068 for *APOE* ε4 homozygous, *p* = 3.65×10^-26^).

Correlation maps of the ten highest-impact features revealed peak associations between feature values and PET voxel intensities predominantly in the temporal lobe (Suppl. Fig. 5), including the anterior and posterior Temporal Fusiform Cortex, Inferior Temporal Gyrus, Temporal Pole, and Planum Polare. Additional clusters of high correlation were observed in the Subcallosal Cortex, Insular Cortex, Frontal Opercular and Frontal Orbital Cortices, and Paracingulate Gyrus (Suppl. Fig. 6). Notably, prominent correlation peaks were also visually identifiable in the cerebellum, which falls outside the coverage of the used Harvard-Oxford cortical brain atlas.

In subsequent analyses, we focused on the high-impact feature set, as it visually represents a denser AD continuum and showed the strongest effect of ranking on Aβ CSF prediction. The results for the core 95% features set are provided in the supplementary materials.

### 3.3. Association of AD continuum clusters with AD-related biomarkers

We performed hierarchical clustering on samples using the selected features and compared the resulting clusters to SUVR-based clusters to assess the agreement between the two approaches (Table 1, Fig. 4A, Suppl. Fig. 7). For the high-impact feature set with two clusters, the two methods agreed on Aβ- and Aβ+ group assignments with concordance of 81.2%. When increasing the number of clusters and excluding samples assigned to the intermediate clusters from analysis, the high-impact feature set achieved its highest agreement of 91.8% for four clusters, with a positive agreement of 92.5% and a negative agreement of 90.4%, indicating well-balanced performance.

**Figure 4.**
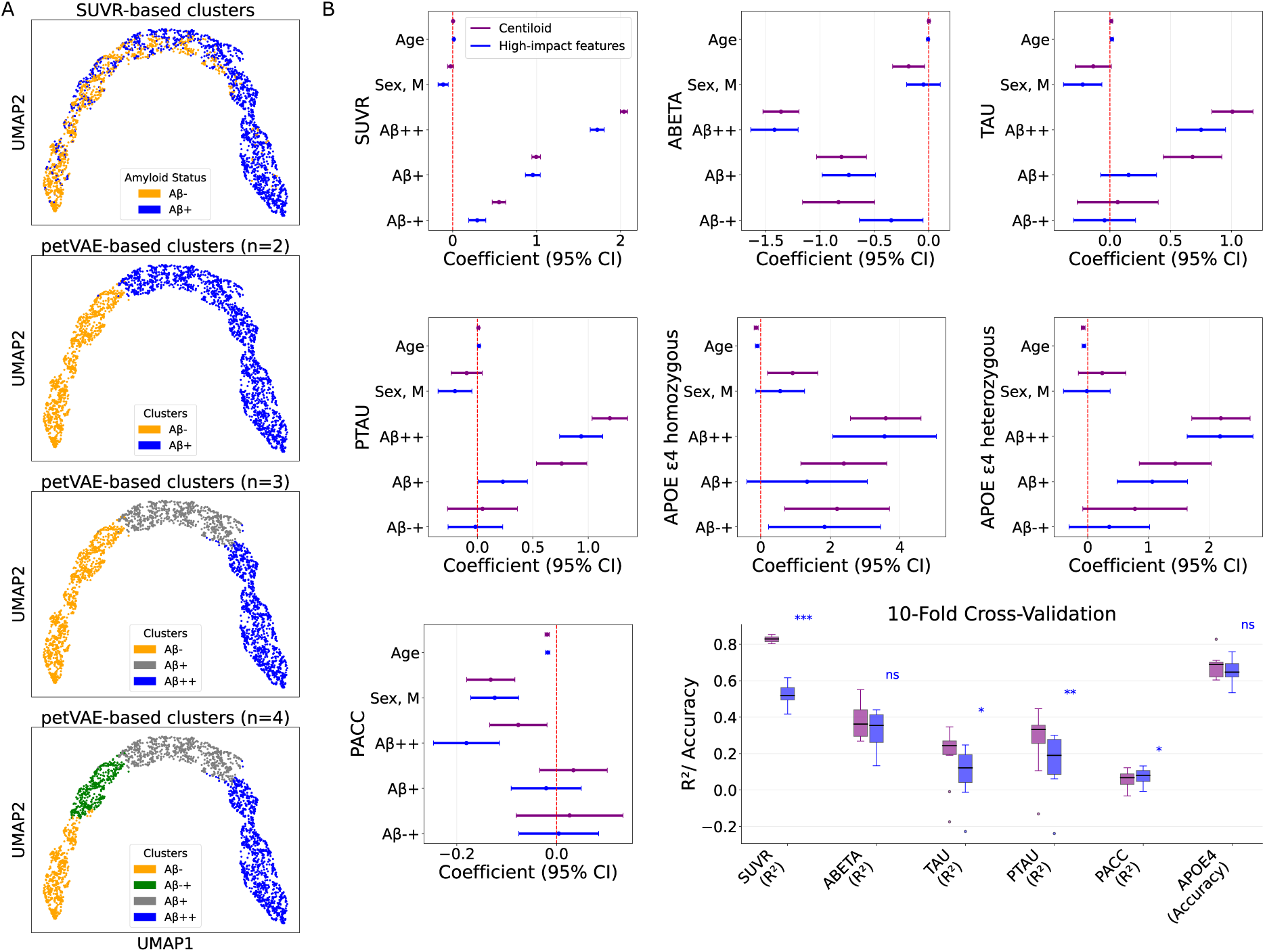
(A) UMAPs embeddings for samples based on high-impact features, colored by SUVR or petVAE-based clusters. (B) Comparison of petVAE- and Centiloid-based clusters by AD-related biomarkers. The Aβ-group was used as the reference group. Abbreviations: SUVR – composite standardized uptake value ratio, ABETA – Aβ cerebrospinal fluid biomarker, TAU – tau cerebrospinal fluid biomarker, PTAU – Phosphorylated tau cerebrospinal fluid biomarker, *APOE* – Apolipoprotein E gene, PACC – Preclinical Alzheimer’s Cognitive Composite score, R^2^-coefficient of determination, ns - not significant, * - p-value< 0.05, ** - p-value< 0.01, *** - p-value< 0.001.

**Table 1.** Performance metrics comparing petVAE-derived hierarchical clusters with SUVR-based Aβ classification. Values are overall agreement/concordance (positive agreement, negative agreement).

| | SUVR (Aβ- & Aβ+) vs petVAE (2 clusters, Aβ- & Aβ+) | SUVR (Aβ- & Aβ+) vs petVAE (3 clusters, Aβ- & Aβ++) | SUVR (Aβ- & Aβ+) vs petVAE (4 clusters, Aβ- & Aβ++) |
| --- | --- | --- | --- |
| Core 95% features | 0.671 (0.469, 0.988) | 0.912 (0.865, 0.980) | 0.912 (0.865, 0.980) |
| High-impact features | 0.812 (0.872, 0.718) | 0.869 (0.823, 0.937) | 0.918 (0.925, 0.904) |

Abbreviations: SUVR - standardized uptake value ratio.

After adjusting to sex and age, Aβ-+ cluster, Aβ+ and Aβ++ were significantly different from Aβ-cluster in SUVR (*p* < 1.64×10^-8^) and CSF Aβ concentration (*p* < 0.02) in baseline samples (Fig. 4B, Suppl. Fig. 9). Cluster Aβ-+ (*n* = 296) had significantly higher odds of being homozygous for the *APOE* ε4 allele (*p* = 0.026) compared to Aβ-cluster (*n* = 378). Cluster Aβ+ (*n* = 511) had significantly higher probability to carry one *APOE* ε4 allele (*p* = 3.36×10^-4^) and higher CSF pTau concentration (*p* = 0.042). Cluster Aβ++ (*n* = 726) is significantly different from Aβ-cluster for all presented AD-related biomarkers (*p* < 3×10⁻⁶).

Centiloid analysis revealed that Aβ-+, Aβ+, and Aβ++ centiloid-based clusters differed from the Aβ-cluster in CSF Aβ levels, SUVR, and likelihood of carrying two APOE ε4 alleles (*p* < 0.004) (Fig. 4B, Suppl. Fig. 9). Compared to the Aβ-cluster, Aβ+ participants showed higher probability of carrying at least one APOE ε4 allele (p < 1.64×10^-8^), and higher CSF tau (p < 1.64×10^-8^) and pTau (p < 1.64×10^-8^) concentrations (Fig. 4B, Suppl. Fig. 9). Poorer cognitive performance was observed only in the Aβ++ cluster compared with Aβ-(*p* = 0.009). In cross-validation analyses, centiloid-based clustering outperformed high-impact feature-based clustering in predicting SUVR (p < 0.001), CSF tau (p < 0.05), and pTau (p < 0.01). In contrast, petVAE-based clustering demonstrated stronger associations with cognitive performance than centiloid-based clustering approach (p < 0.05). No significant differences were observed between the two clustering approaches in Aβ CSF prediction.

### 3.4. Survival analysis

We evaluated differences in AD progression among the identified clusters using survival analysis. Distinct survival trajectories were observed among clusters, suggesting variable risk of progression during 72 months. The log-rank test indicated a statistically significant difference in survival distributions between clusters (*p* = 8.61×10^-19^). Clusters Aβ- and Aβ-+ showed the slowest rate of progression, cluster Aβ++ exhibited the most rapid decline toward AD diagnosis, while cluster Aβ+ performed a moderate rate of AD progression (Fig. 5A, Suppl. Fig. 10).

**Figure 5.**
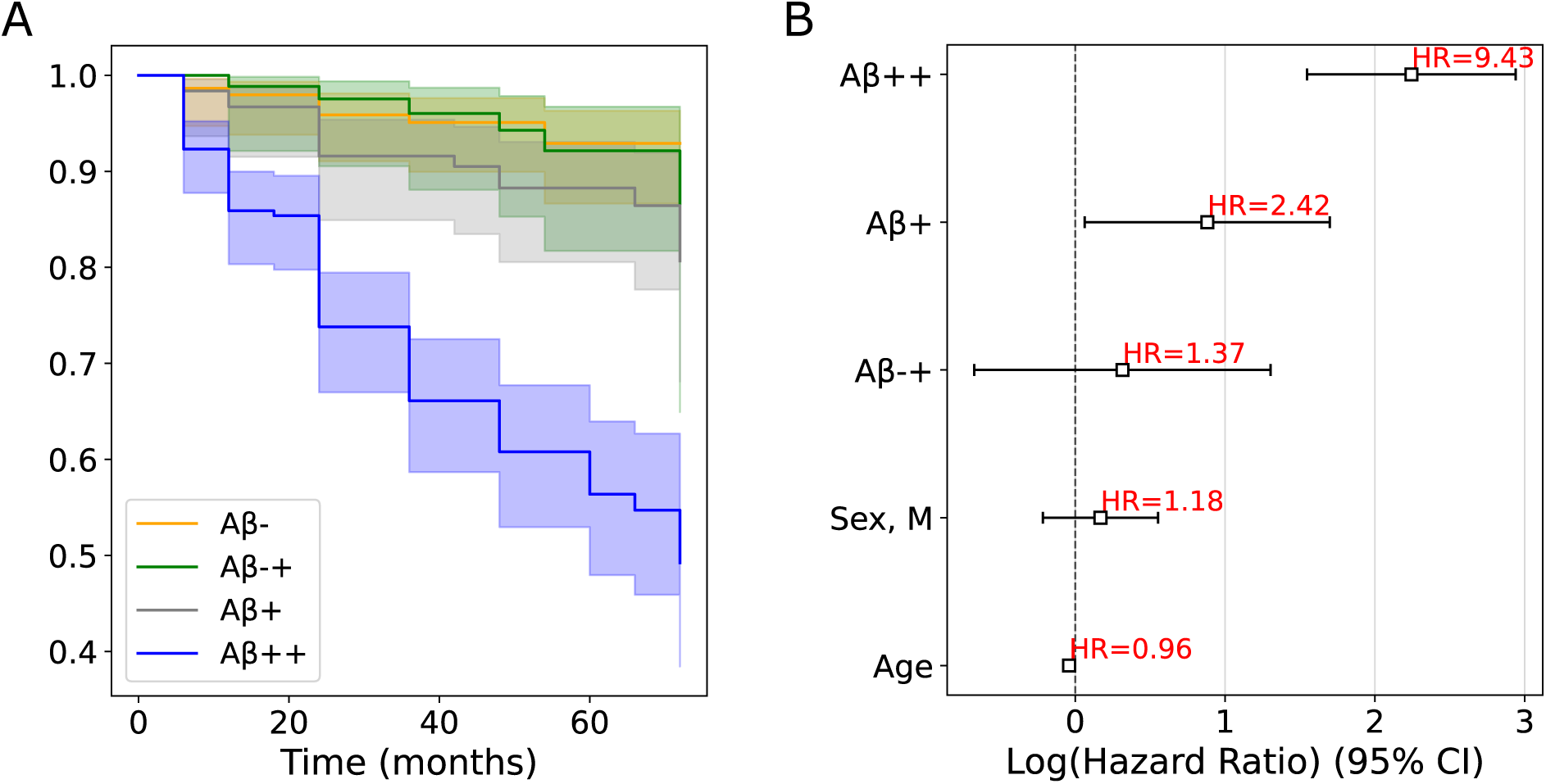
Survival analysis of petVAE-based clusters. (A) Kaplan–Meier estimated survival curves with 95% confidence intervals. (B) Comparison of clusters by Cox proportional hazards models reporting hazard ratios with 95% confidence intervals with age, sex as covariates.

To quantify these differences, we fitted Cox proportional hazards models with cluster membership as the primary predictor, adjusting for age and sex (Fig. 5B, Suppl. Fig. 10). Compared with the reference cluster (Aβ-), participants in the Aβ-+ cluster did not show a significantly different risk of progression to AD (HR = 1.37, *p* = 0.55). Clusters Aβ+ and Aβ++ showed elevated hazards (Aβ+ HR = 2.42, *p* = 1.17×10^-7^; Aβ++ HR = 9.43, *p* = 1.31×10^-15^), indicating that individuals in these groups experienced a faster rate of AD progression than those in the Aβ-group.

### 3.5. Translation to other tracers

Despite differences among PET scans acquired using AV45, PiB, and FBB Aβ tracers, petVAE reconstructed all images with good accuracy (Fig. 6A–B). As expected, reconstruction performance was highest for AV45, which was the tracer used to train and validate the model (μSSIM = 0.90, μPSNR = 29.6 dB). The model also reconstructed PiB scans with an average SSIM of 0.84 and PSNR of 27.8 dB, and FBB scans with an average SSIM of 0.84 and PSNR of 27.3 dB. After feature selection and UMAP embedding, AV45 and FBB features showed stronger correspondence and distributed more evenly along the continuum, whereas PiB features clustered tightly at one end and formed a small, distinct group (Fig. 6B).

**Figure 6.**
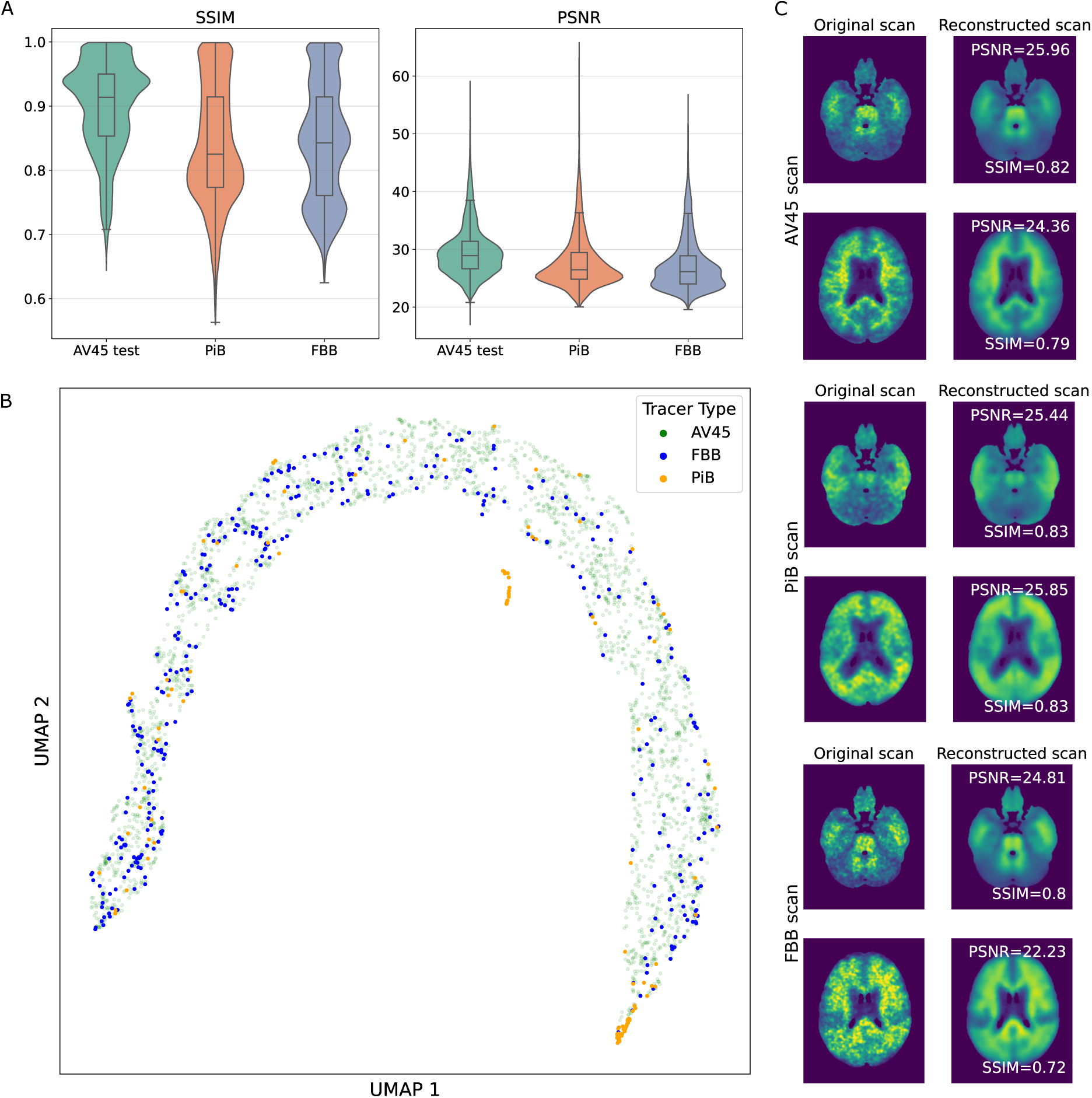
Evaluation of petVAE ability to reconstruct PET scans obtained with PiB and FBB tracers. (A) Distribution of SSIM and PSNR values for AV45, PiB and FBB Aβ PET scans. (B) UMAP embedding for AV45, PiB and FBB Aβ PET scans based on features with an importance score greater than 0.01, colored by Aβ tracer. (C) Example of AV45, PiB and FBB Aβ PET slices and their reconstruction by the petVAE. Abbreviations: SSIM - Structural Similarity Index Measure, PSNR - Peak Signal-to-Noise Ratio, AV45 - [^18^F]-Florbetapir Aβ PET tracer, [^11^C]-PiB - Pittsburgh compound B Aβ PET tracer, FBB - [^18^F]-Florbetaben Aβ PET tracer.

## 4. Discussion

Based on petVAE reconstruction and clustering of PET scans, we were able to go beyond conventional binary classification of Aβ negativity versus positivity and detected four subgroups along the AD continuum. These included two extreme Aβ groups (Aβ-, Aβ++), corresponding to the conventional Aβ negative and Aβ positive categories, as well as two intermediate groups (Aβ-+, Aβ+). The intermediate subgroups showed divergence from the lowest Aβ group (Aβ-) in CSF and SUVR measures of Aβ accumulation, as well as in their genetic risk for AD. Although cognitive performance differed between the extreme Aβ groups, neither intermediate group exhibited measurable cognitive impairment relative to the Aβ-group. Nevertheless, the intermediate group positioned further along the AD continuum (Aβ+) demonstrated a faster rate of progression to AD. Therefore the intermediate subgroups represent individuals at elevated risk for AD development despite the absence of pronounced clinical symptoms. Defining and characterizing such intermediate subgroups is crucial for enabling earlier diagnosis and informing more effective treatment strategies in AD.

To our knowledge, petVAE is the first VAE-based model applied to Aβ PET to delineate not only Aβ negative and Aβ positive groups, but also intermediate subgroups along the Aβ continuum. Several previous studies have used data-driven approaches to identify stages of AD progression or to characterize subgroups along the tau-aggregation continuum [20,41]. A recent study utilized a VAE model to identify four stages of tau aggregation; however, only one stage differed from the others in cognitive performance, tau, and amyloid aggregation [41]. This highlights the importance of using Aβ scans, CSF or plasma biomarkers for earlier detection of AD development. In our study, using only Aβ PET scans, the latent space learned by petVAE captured the continuum of Aβ aggregation and enabled the identification of four distinct groups differing in Aβ burden, pTau levels, and cognitive performance.

The four sub-groups identified by petVAE-based clustering reflected a progression pattern in line with the amyloid cascade theory. While the differences between the Aβ- and Aβ++ groups were evident across SUVR values, all CSF AD-biomarkers, cognitive scores, genetic risk and rate of AD progression, we also observed significant variation among the intermediate clusters when compared against the Aβ-group. Based on survival analysis, the Aβ-+ group most closely resembled the Aβ-samples; although Aβ aggregation levels were significantly higher (SUVR and CSF Aβ), the rate of AD progression was not increased. However, the Aβ-+ group had higher odds of being *APOE* ε4 homozygous. Stronger separation was observed for Aβ+ samples where we already saw significantly higher concentration of pTau in CSF but not yet the worse cognitive performance. Thus, the Aβ-+ group showed significantly higher Aβ; the Aβ+ group exhibited significantly higher Aβ and pTau; and the Aβ++ group additionally showed significant differences in cognition. This finding aligns with cascade theory whereby amyloid aggregation occurs before tau accumulation which eventually causes cognitive decline [42]. According to Cox proportional hazards models, Aβ+ also had an almost 2.5-fold higher hazard of progressing to AD compared with Aβ-.

In comparison to the conventional Centiloid grouping, petVAE-based clustering showed stronger associations with cognitive performance, indicating that our data-driven representations of PET signal captured clinical symptoms as measured with PACC score that has been developed to assess beta amyloid related cognitive change [29]. This difference may reflect the ability of the deep learning approach to incorporate more complex, spatially distributed patterns of tracer uptake beyond global burden measures [43,44]. No significant differences were observed between clustering approaches in predicting CSF Aβ, suggesting that both methods capture core Aβ-related pathology to a comparable extent. Centiloid-based clustering was more effectively capturing variability in SUVR, tau, and pTau. Because Centiloids were derived from SUVR itself, their stronger association with this measure was expected. The stronger performance for CSF tau and pTau may reflect the well-established relationship between global Aβ burden and downstream tau phosphorylation. This is consistent with evidence that CSF pTau phosphorylation occupancies correlate strongly and continuously with amyloid PET Centiloid values [45,46] petVAE-derived clusters - based on spatial uptake patterns rather than global magnitude - may be less directly associated with tau pathology. Together, these results highlight that centiloid- and petVAE-based clustering approaches may provide complementary information, with centiloid clustering more closely aligned with established AD biomarker measures and petVAE-derived clusters potentially offering additional sensitivity to cognitive markers of AD.

Based on petVAE reconstructions of PiB and FBB Aβ tracers, the model achieved reconstruction quality comparable to that of AV45 tracer scans, on which it was trained. However, in the AD continuum representation, we observed a very noticeable separation of PiB scans, whereas FBB scans integrated well. This can be explained by the differences between ¹¹C-labelled (PiB) and ¹⁸F-labelled (FBB and AV45) amyloid tracers [47,48]. PiB was the first Aβ tracer developed for use in humans and is often considered the gold-standard research tracer [49,50]. It exhibits higher cortical binding and higher contrast, and lower non-specific white-matter uptake compared with ¹⁸F-labelled tracers. However, due to its very short half-life and the need for on-site production, PiB is not widely used in clinical practice. We hypothesize that although our model is capable of reconstructing PiB scans, the latent features captured the strong differences between PiB and the ¹⁸F-labelled tracers, resulting in the pronounced separation observed along the AD continuum.

The petVAE model may provide some robustness across PET scans acquired with different ¹⁸F-labelled amyloid tracers and different scanners, suggesting reduced sensitivity to common sources of technical variability. In particular, petVAE applies a level of smoothing to the reconstructed scans that is comparable to that used in conventional PET image-processing pipelines, which helps reduce scanner- and tracer-related differences before voxel-wise statistical testing [51]. Moreover, petVAE’s fully data-driven design enables automated extraction of latent imaging features and unbiased identification of AD-related subgroups without relying on predefined regions of interest or SUVR thresholds, representing an important advantage over conventional PET quantification methods [52]. In clinical practice, Aβ PET classification is rarely performed in a data-driven manner directly from the scans. Instead, scans are typically interpreted manually by certified readers [47] or categorized using quantification metrics such as SUVR, Centiloid scale, Z-scores, or Aβ load [10,53,54]. These approaches are heavily dependent on study-specific or protocol-specific thresholds, which limits scalability and complicates clinical implementation [52]. By contrast, the petVAE framework provides a continuous, standardized, and threshold-free representation of amyloid burden, offering a more flexible and potentially more clinically generalizable alternative.

Moving forward, the model requires further validation on fully independent external datasets, including prospective clinical data, before any clinical translation can be considered. Although the model showed preliminary indications of robustness across different amyloid tracers, this observation remains largely qualitative and requires validation in larger, tracer-diverse datasets, as well as through dedicated quantitative analyses. Another important topic for further investigation is the role of feature importance within the model. If high-impact features are sufficient for defining the Aβ continuum, it is necessary to clarify the contribution of the remaining features. In particular, it should be assessed whether excluding these lower-impact features would affect reconstruction quality, and whether the model could be simplified without compromising performance. While preliminary voxel-wise correlation analyses of the ten highest-impact features revealed spatial patterns of association predominantly in the temporal lobe, limbic, and prefrontal regions, areas known to be implicated in early amyloid deposition [55], a dedicated follow-up study is required to strengthen the anatomical interpretation of these findings and to extend this analysis to the full core 95% feature set. Finally, additional development is needed to integrate petVAE into a streamlined, reproducible pipeline (*e.g.*, using workflow managers such as Nextflow) to facilitate broader adoption and routine use.

In summary, petVAE provided a fully data-driven framework for reconstructing Aβ PET scans and identifying subgroups along the AD continuum, including previously uncharacterized intermediate subgroups. The four subgroups detected by petVAE align with the amyloid cascade theory and show differentiated profiles in Aβ, pTau, cognitive performance, and genetic risk, highlighting individuals at elevated risk for AD despite minimal clinical symptoms.

The model demonstrates robustness across different ¹⁸F-labelled tracers and scanners, and offers a continuous, threshold-free representation of amyloid burden that addresses limitations of conventional binary PET classification. While further validation on independent datasets and comparison with standardized metrics are necessary, petVAE represents a promising approach for early detection, risk stratification, and potentially guiding clinical decision-making in AD.

## Acknowledgement

The authors wish to acknowledge CSC – IT Center for Science, Finland, for generous computational resources.

This study was supported by the Doctoral Programme in Population Health of the University of Helsinki (AT), Finnish Brain Foundation (20250101 to AT), Yrjö Jahnsson Foundation (20257996 to AT) and the Research Council of Finland (322675 and 328890 to EP). EV was supported by the Sigrid Jusélius Foundation.

Data collection and sharing for the Alzheimer’s Disease Neuroimaging Initiative (ADNI) is funded by the National Institute on Aging (National Institutes of Health Grant U19AG024904). The grantee organization is the Northern California Institute for Research and Education. In the past, ADNI has also received funding from the National Institute of Biomedical Imaging and Bioengineering, the Canadian Institutes of Health Research, and private sector contributions through the Foundation for the National Institutes of Health (FNIH) including generous contributions from the following: AbbVie, Alzheimer’s Association; Alzheimer’s Drug Discovery Foundation; Araclon Biotech; BioClinica, Inc.; Biogen; Bristol-Myers Squibb Company; CereSpir, Inc.; Cogstate; Eisai Inc.; Elan Pharmaceuticals, Inc.; Eli Lilly and Company; EuroImmun; F. Hoffmann-La Roche Ltd and its affiliated company Genentech, Inc.; Fujirebio; GE Healthcare; IXICO Ltd.; Janssen Alzheimer Immunotherapy Research & Development, LLC.; Johnson & Johnson Pharmaceutical Research & Development LLC.; Lumosity; Lundbeck; Merck & Co., Inc.; Meso Scale Diagnostics, LLC.; NeuroRx Research; Neurotrack Technologies; Novartis Pharmaceuticals Corporation; Pfizer Inc.; Piramal Imaging; Servier; Takeda Pharmaceutical Company; and Transition Therapeutics.The A4 Study is a secondary prevention trial in preclinical Alzheimer’s disease, aiming to slow cognitive decline associated with brain amyloid accumulation in clinically normal older individuals. The A4 Study is funded by a public-private-philanthropic partnership, including funding from the National Institutes of Health-National Institute on Aging, Eli Lilly and Company, Alzheimer’s Association, Accelerating Medicines Partnership, GHR Foundation, an anonymous foundation and additional private donors, with in-kind support from Avid and Cogstate. The companion observational Longitudinal Evaluation of Amyloid Risk and Neurodegeneration (LEARN) Study is funded by the Alzheimer’s Association and GHR Foundation. The A4 and LEARN Studies are led by Dr. Reisa Sperling at Brigham and Women’s Hospital, Harvard Medical School and Dr. Paul Aisen at the Alzheimer’s Therapeutic Research Institute (ATRI), University of Southern California. The A4 and LEARN Studies are coordinated by ATRI at the University of Southern California, and the data are made available through the Laboratory for Neuro Imaging at the University of Southern California. The participants screening for the A4 Study provided permission to share their de-identified data in order to advance the quest to find a successful treatment for Alzheimer’s disease. We would like to acknowledge the dedication of all the participants, the site personnel, and all of the partnership team members who continue to make the A4 and LEARN Studies possible. The complete A4 Study Team list is available on: a4study.org/a4-study-team.

## Ethics and Integrity Statement

This study used existing data from the Alzheimer’s Disease Neuroimaging Initiative (ADNI) and the Anti-Amyloid Treatment in Asymptomatic Alzheimer’s Disease (A4) study. All required dataset acknowledgements and citations are included in the manuscript, along with full funding acknowledgements.

The authors declare no conflicts of interest.

As this research relied exclusively on previously collected, de-identified data from established research cohorts, no additional ethics approval or patient consent was required for this study. No material from other sources requiring reproduction permission was used. This work does not involve a clinical trial and therefore does not require clinical trial registration.

## Notes

### Competing Interest Statement

The authors have declared no competing interest.

### Summary of Updates

All sections of the manuscript were improved and updates according to reviewers' comments.

