## Supplementary file for "petVAE: A Data-Driven Model for Identifying Amyloid PET Subgroups Across the Alzheimer’s Disease Continuum"

Supplementary methods

Visualization of brain areas represented by petVAE features

To visualize high-impact features and brain areas that they represent, we computed Pearson correlation maps. Each map reflects the voxel-wise association between a single feature value and PET signal intensity across all scans in the dataset. Higher absolute correlation values indicate brain regions where the feature most strongly covaries with tracer uptake. Correlation maps were computed in native image space and visualized on the MNI152 standard brain template. For each feature, the axial slice with the highest peak correlation was selected for display. We used the Harvard-Oxford cortical atlas (Makris et al. 2006) to extract mean correlation values per anatomical region and sort regions by their association strength. Due to the large number of features in the core 95% feature set (n = 3,098), voxel-wise correlation mapping was not performed for it.

Supplementary Table 1. Population characteristics of the dataset

| | ADNI | A4 |
|---|---|---|
| Number of subjects | 978 | 1,328 |
| Number of PET scans | 1,782 | 1,328 |
| Age (mean±sd) | 74.9±7.4 | 71.8±4.8 |
| Sex (female) | 816 (45.8%) | 790 (59.5%) |
| Diagnosis:<br>CN<br>SMC<br>MCI<br>AD | <br>495 (27.8%)<br>163 (9.1%)<br>967 (54.3%)<br>157 (8.8%) | <br>-<br>-<br>-<br>- |
| CDR:<br>0.0<br>0.5<br>1.0<br>2.0<br>3.0 | n = 1762<br>645 (36.6%)<br>915 (51.9%)<br>178 (10.1%)<br>18 (1.0%)<br>6 (0.3%) | <br>1328 (100%)<br>-<br>-<br>-<br>- |
| Aβ status SUVR-based:<br>Aβ-<br>Aβ+ | n = 1,624<br>785 (48.3%)<br>839 (51.7%) | n = 1,328<br>320 (24.1%)<br>1,008 (75.9%) |
| PACC score (mean±sd) | -4.6±7.0 | -0.1±1.0 |
| Number of Apolipoprotein E ε4-alleles:<br>0<br>1<br>2 | n = 1,776<br><br>1,024 (57.7%)<br>609 (34.3%)<br>143 (8.0%) | <br><br>-<br>-<br>- |

Abbreviations: CN – cognitively normal, SMC – subjective memory complaints, MCI - mild cognitive impairment, AD – Alzheimer’s disease, CDR - clinical dementia rating, Aβ – Amyloid-β , PACC – preclinical Alzheimer cognitive composite, SUVR – standardized uptake value ratio, PET – positron emission tomography, ADNI – Alzheimer’s Disease Neuroimaging Initiative, A4 – Anti-Amyloid Treatment in Asymptomatic Alzheimer's Diseas

Supplementary Table 2. Population characteristics of petVAE feature-based clusters

| | | | Mean SUVR ± SD (N=2802) | Aβ+ (N=1713, 61.1%) | Aβ CSF (N=760) | pTau CSF (N=961) | PACC (N=2800) | APOE ε4-carrier (N=1472) | Mean age ± SD (N=2802) | Sex (Female, N=1470, 52.5%) |
|---|---|---|---|---|---|---|---|---|---|---|
| Core 95% features | 3 Clusters | Aβ- | 1.03 ± 0.1 | 125, 16.7% of samples in the cluster | 1150.7 ± 331.3 | 19.4 ± 7.3 | -1.4 ± 3.8 | 144, 21.7 % of samples in the cluster | 74.45 ± 7.1 | 251, 33.5% of samples in the cluster |
| | | Aβ+ | 1.18 ± 0.2 | 784, 63.5% | 993.9 ± 350.2 | 27.1 ± 16.2 | -0.4 ± 2.9 | 119, 38.4% | 71.19 ± 5.4 | 847, 68.6% |
| | | Aβ++ | 1.45 ± 0.2 | 804, 98.4 % | 689.9 ± 219.4 | 34.1 ± 14.3 | -4.5 ± 6.8 | 321, 64.6% | 75.49 ± 6.1 | 372, 45.5% |
| | 4 Clusters | Aβ- | 1.0 ± 0.1 | 125,16.7% | 1150.7 ± 331.3 | 19.4 ± 7.3 | -1.4 ± 3.8 | 144, 21.7% | 74.45 ± 7.1 | 251, 33.5% |
| | | Aβ-+ | 1.1 ± 0.1 | 251,37.6% | 1112.7 ± 350.7 | 21.9 ± 9.3 | -0.3 ± 2.7 | 77, 32.5% | 71.0 ± 5.7 | 482, 72.2% |
| | | Aβ+ | 1.3 ± 0.2 | 533,94.0% | 821.1 ± 270.3 | 39.6 ± 21.8 | -0.6 ± 3.0 | 42, 57.5% | 71.5 ± 5.1 | 365, 64.4% |
| | | Aβ++ | 1.5 ± 0.2 | 804,98.4% | 689.9 ± 219.4 | 34.1 ± 14.3 | -4.5 ± 6.8 | 321, 64.6% | 75.5 ± 6.1 | 372, 45.5% |

| | | | | | | | | | | |
|---|---|---|---|---|---|---|---|---|---|---|
| High-impact features | 3 Clusters | Aβ- | 1.0 ± 0.1 | 219, 21.9% | 1241.8 ± 310.5 | 19.9 ± 7.2 | -0.7 ± 3.3 | 110, 19.3% | 72.0 ± 6.3 | 538, 53.8% |
| | | Aβ+ | 1.2 ± 0.2 | 477, 65.3% | 948.1 ± 317.5 | 24.9 ± 13.9 | -1.4 ± 3.9 | 133, 38.7% | 73.7 ± 6.7 | 410, 56.1% |
| | | Aβ++ | 1.4 ± 0.2 | 1017, 95.1% | 722.5 ± 256.1 | 33.1 ± 15.5 | -3.4 ± 6.1 | 341, 61.2% | 74.3 ± 6.1 | 522, 48.8% |
| | 4 Clusters | Aβ- | 1.0 ± 0.1 | 83, 14.3% | 1300.7 ± 284.7 | 19.4 ± 6.3 | -0.7 ± 3.5 | 50, 14.0% | 72.3 ± 6.5 | 315, 54.1% |
| | | Aβ-+ | 1.1 ± 0.1 | 136, 32.5% | 1154.2 ± 328.0 | 20.8 ± 8.6 | -0.8 ± 3.2 | 60, 28.0% | 71.5 ± 6.1 | 223, 53.2% |
| | | Aβ+ | 1.2 ± 0.2 | 477, 65.3% | 948.1 ± 317.5 | 24.9 ± 13.9 | -1.4 ± 3.9 | 133, 38.7% | 73.7 ± 6.7 | 410, 56.1% |
| | | Aβ++ | 1.4 ± 0.2 | 1017, 95.1% | 722.5 ± 256.1 | 33.1 ± 15.5 | -3.4 ± 6.1 | 341,61.2% | 74.3 ± 6.1 | 522, 48.8% |

Abbreviations: Aβ – Amyloid-β, pTau - phosphorylated tau, PACC – preclinical Alzheimer cognitive composite, SUVR – standardized uptake value ratio, CSF - cerebrospinal fluid, APOE – Apolipoprotein E gene, SD - standard deviation, N - number of available observations for variable, Aβ-/Aβ-+/Aβ+/Aβ++ denote petVAE-based hierarchical clustering–defined groups representing increasing levels of Aβ load on PET imaging and corresponding positions along the Alzheimer's disease continuum.

Supplementary Table 3. High-impact features, correspondent PET slices and positions at original 64-dimentional latent vector

| Feature ID | PET scan slice index | Location at latent vector |
|---|---|---|
| 2905 | 45 | 25 |
| 3014 | 47 | 6 |
| 3033 | 47 | 25 |
| 3161 | 49 | 25 |
| 3225 | 50 | 25 |
| 3270 | 51 | 6 |
| 3277 | 51 | 13 |
| 7022 | 109 | 46 |
| 7086 | 110 | 46 |
| 7693 | 120 | 13 |

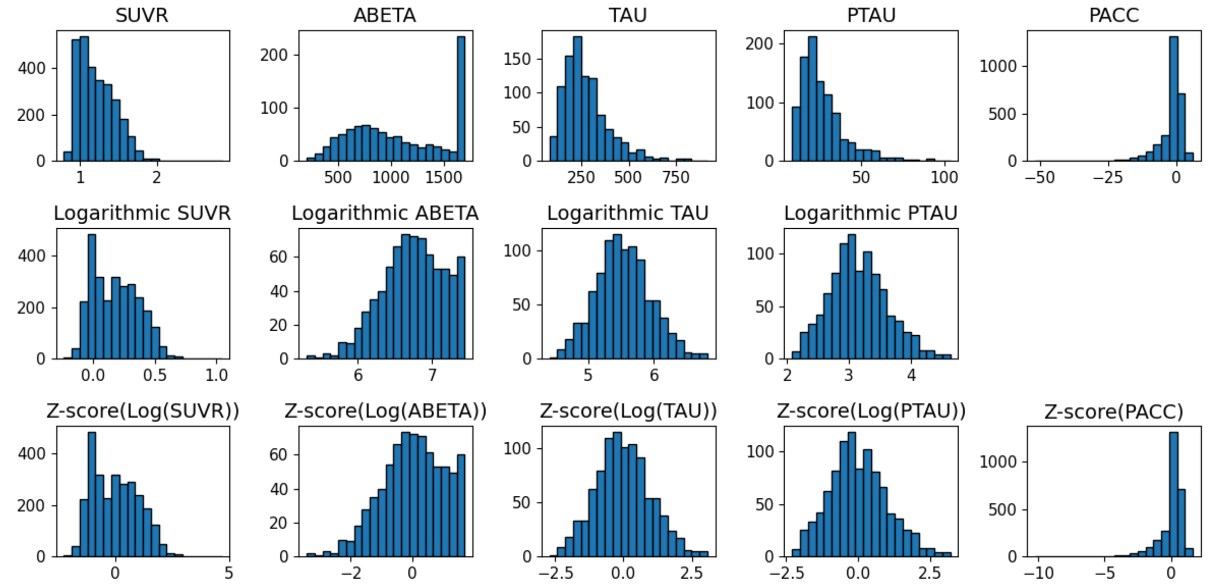

Supplementary Figure 1. Distribution of original values of composite SUVR, CSF biomarker measurements (ABETA, TAU and PTAU), and PACC scores (top row), after logarithmic transformation (middle row) and after z-transformation (bottom row).

Abbreviations: SUVR – composite standardized uptake value ratio, ABETA – Aβ cerebrospinal fluid biomarker, PACC – Preclinical Alzheimer's Cognitive Composite score, TAU – tau cerebrospinal fluid biomarker, PTAU – phosphorylated tau cerebrospinal fluid biomarker

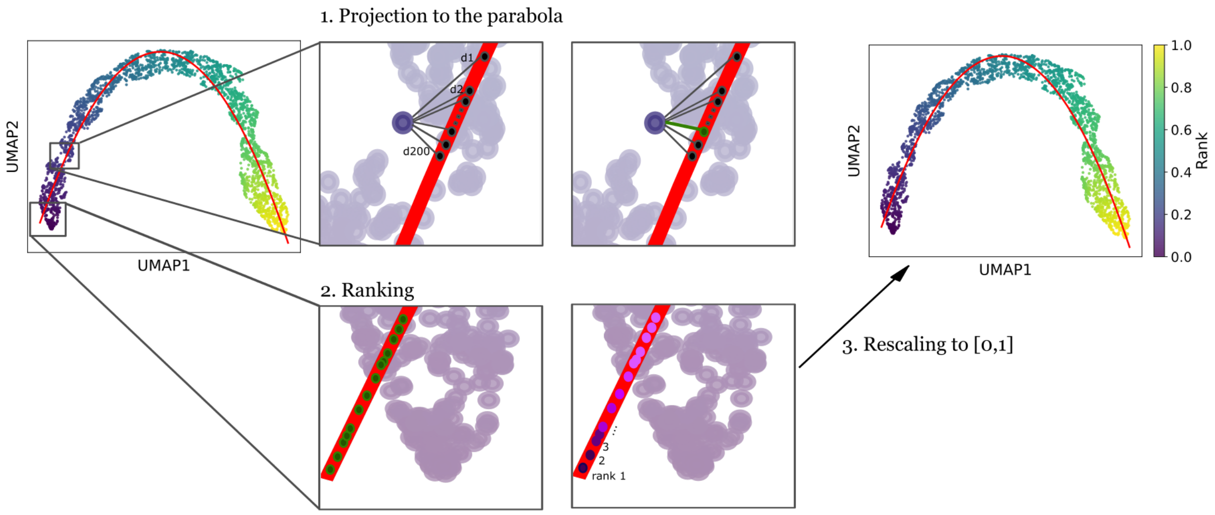

Supplementary Figure 2. Schematic illustration of PET scan ranking in three steps. In the first step, we generated 200 candidate points on the parabola within a small interval around each original point and selected the candidate closest to the original point as its projection. In the second step, the projections were sorted and ranked according to their position along the parabola. Finally, the ranks were normalized to the interval [0,1]. Abbreviations: UMAP - Uniform Manifold Approximation and Projection; d1, d2, …, d200 - Euclidean distance between the original point and each candidate point.

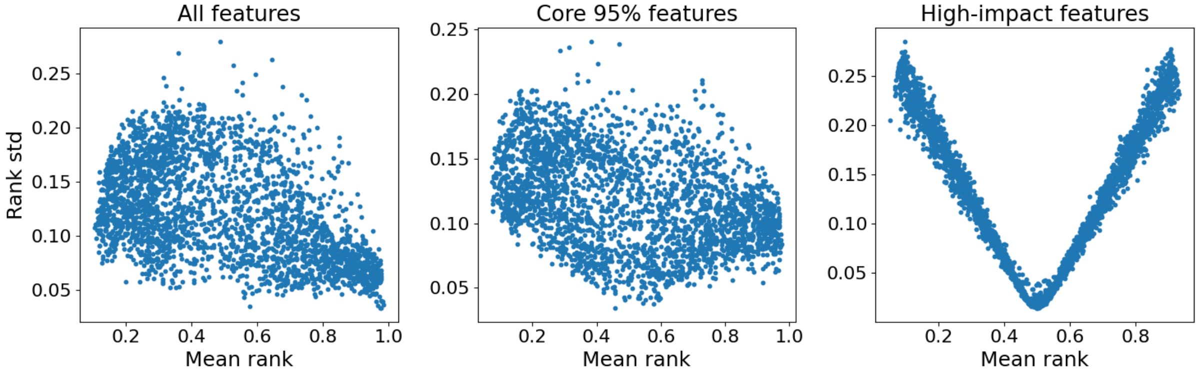

Supplementary Figure 3. Stability of PET scan rankings along the Alzheimer's disease continuum derived from UMAP embeddings. Each point represents a scan, positioned by its mean rank (x-axis) and rank standard deviation (y-axis), computed across 1,000 UMAP bootstrap resamples and subsequent ranking procedures. Lower standard deviation (std) indicates more stable (consistent) placement along the trajectory, whereas higher values reflect increased ranking uncertainty. The three panels show results for all features (left), the core 95% of features (middle), and high-impact features (right), highlighting how feature selection influences ranking stability across the continuum.

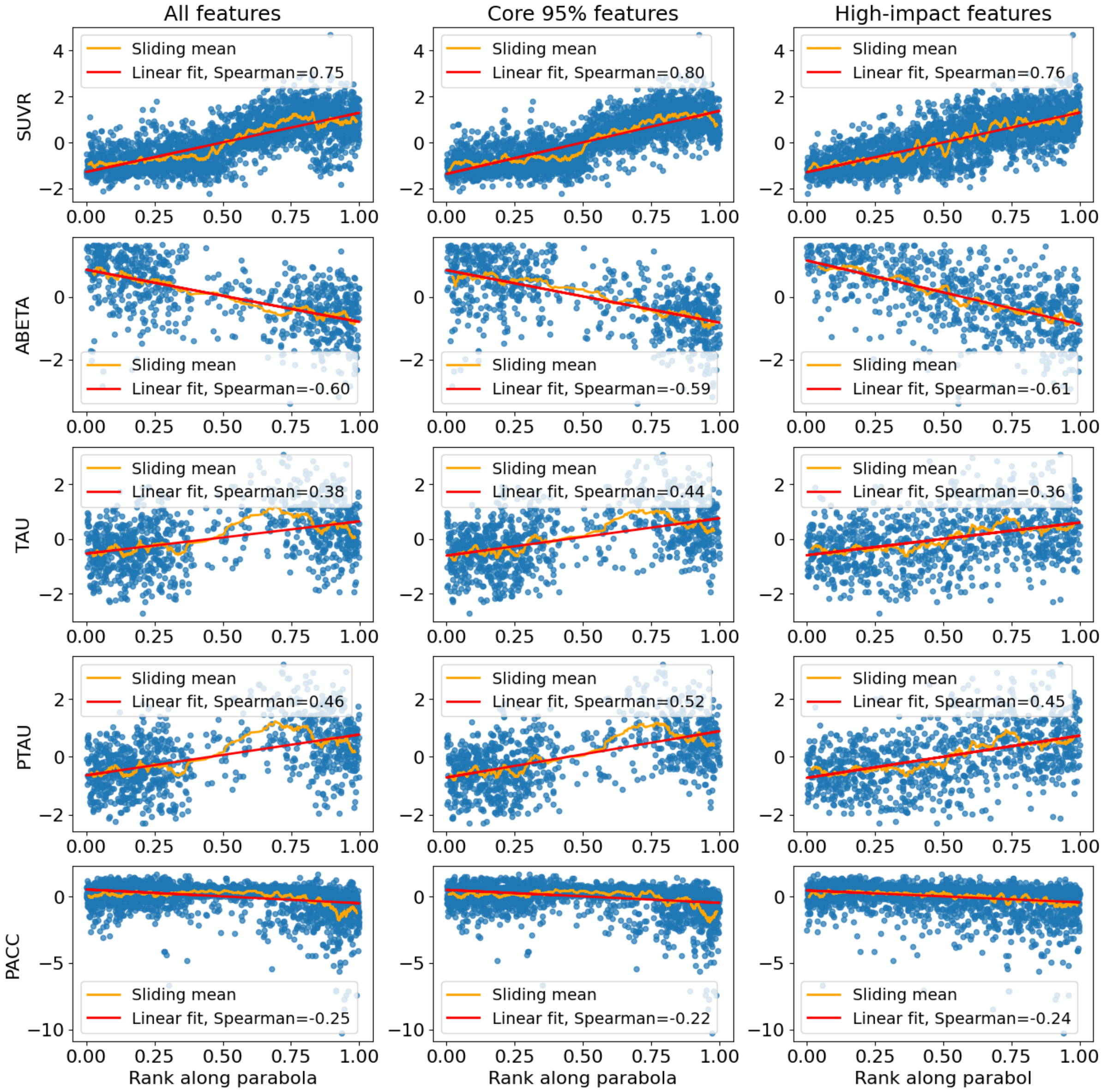

Supplementary Figure 4. Correlation between assigned ranks and AD-related biomarkers. Abbreviations: SUVR – composite standardized uptake value ratio, ABETA – Aβ cerebrospinal fluid biomarker, PACC – Preclinical Alzheimer's Cognitive Composite score, TAU – tau cerebrospinal fluid biomarker, PTAU – Phosphorylated tau cerebrospinal fluid biomarker.

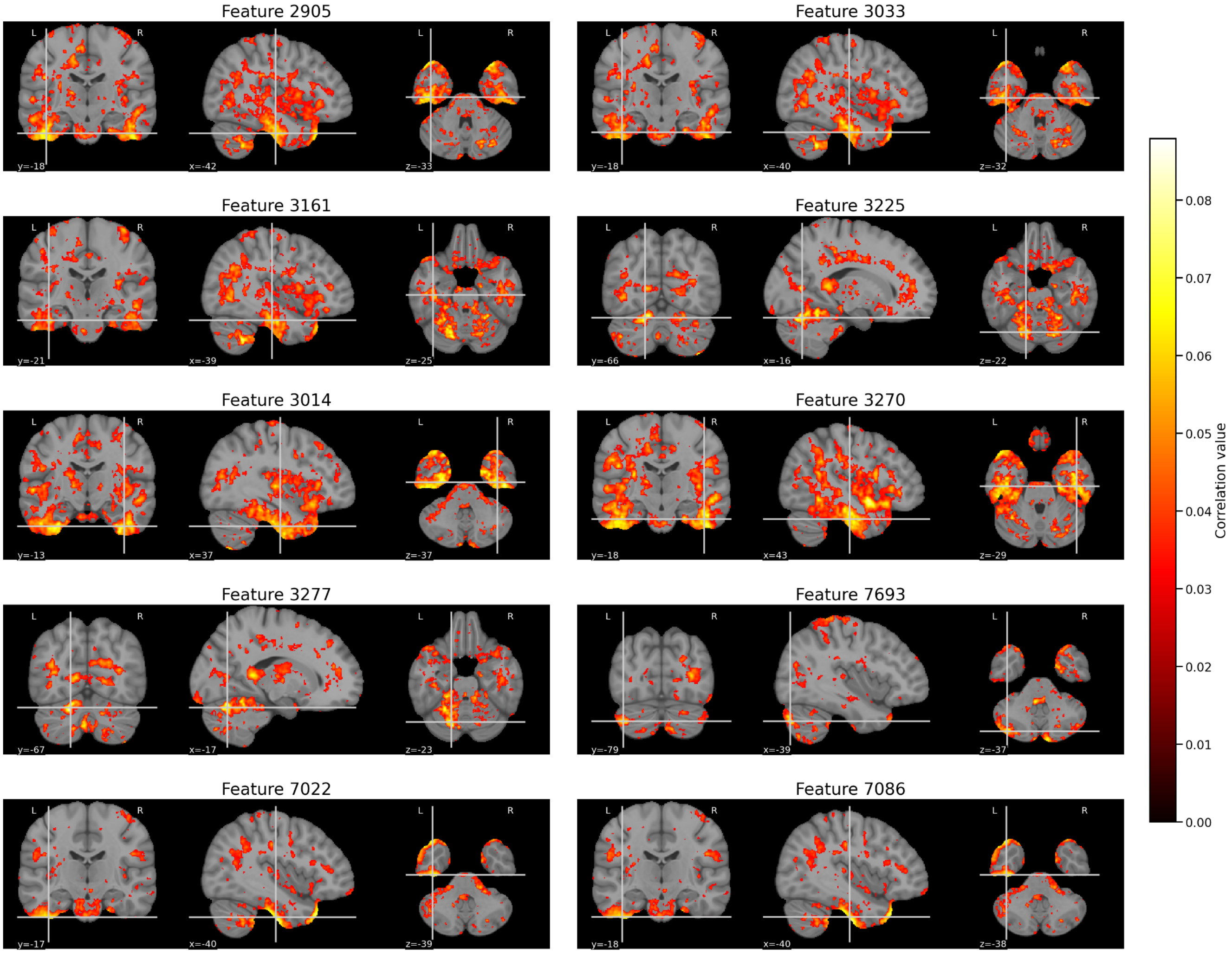

Supplementary Figure 5. Pearson correlation maps between the high-impact features and PET voxel intensity across all scans (n = 3,110). Maps display the absolute Pearson correlation coefficient at each voxel, thresholded at |r| > 0.03 to highlight regions of strongest association. Visualizations are centered at the peak correlation coordinate for each feature and overlaid on the MNI152 standard brain template.

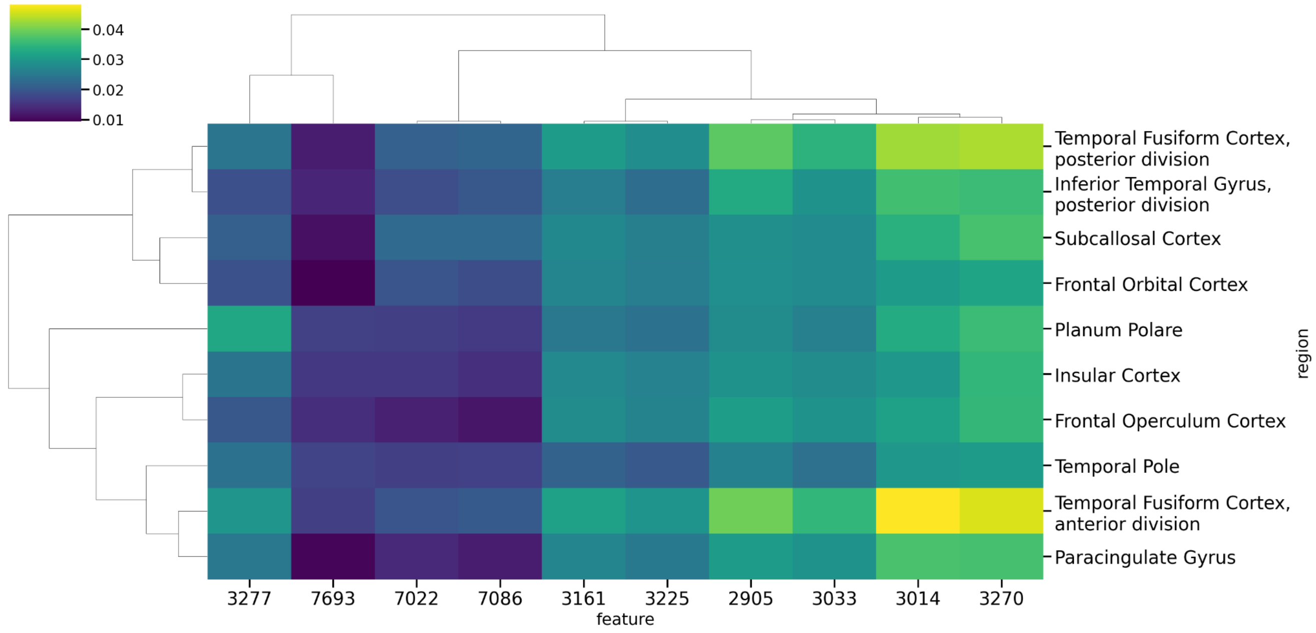

Supplementary Figure 6. Mean absolute Pearson correlation between the high-impact features and PET signal intensity across ten top-ranked cortical regions. Regions were selected based on the highest average correlation across all ten features. Hierarchical clustering was applied to both features and regions. Color scale represents the mean absolute correlation coefficient per feature-region pair.

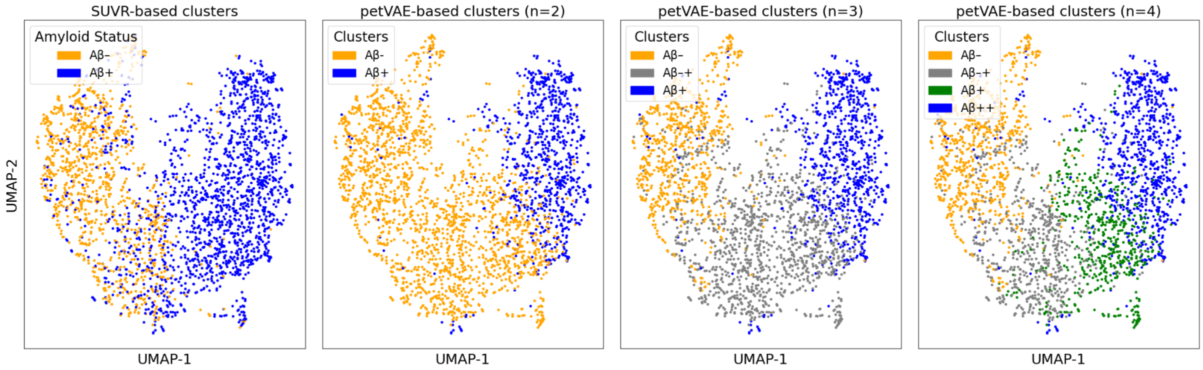

Supplementary Figure 7. UMAPs embeddings for samples based on features that cover 95% of Random Forest total importance, colored by SUVR (first plot from the left) and petVAE-based clusters (plots 2-4 from the left). Abbreviations: SUVR – standardized uptake value ratio, Aβ – Amyloid-β, Aβ-/Aβ-+/Aβ+/Aβ++ denote petVAE-based hierarchical clustering–defined groups representing increasing levels of Aβ load on PET imaging and corresponding positions along the Alzheimer's disease continuum.

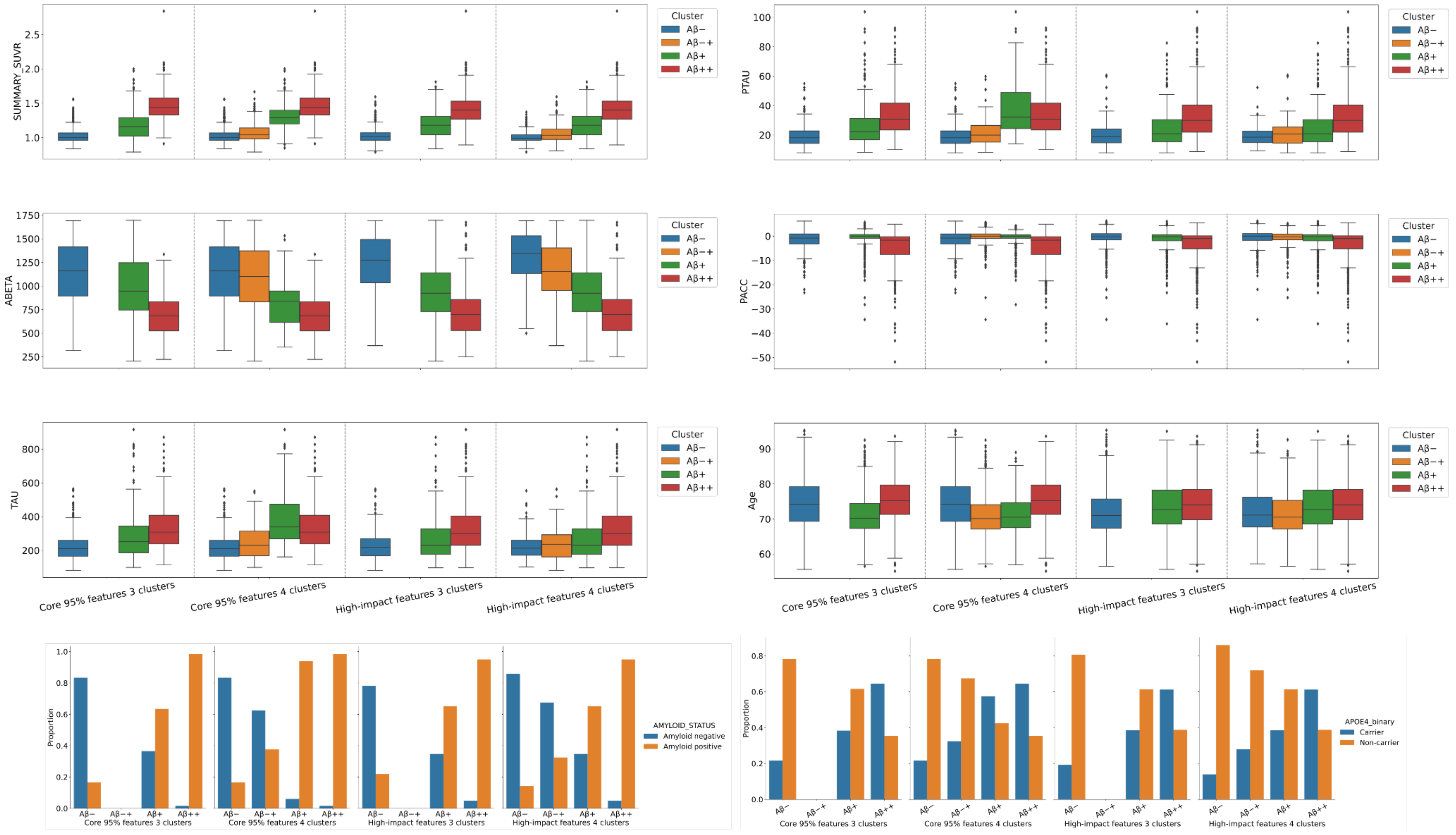
SUMMARY_SUVR
PTAU
ABETA
PACC
TAU
Age
Cluster
Aβ−
Aβ−+
Aβ+
Aβ++
Core 95% features 3 clusters
Core 95% features 4 clusters
High-impact features 3 clusters
High-impact features 4 clusters
Proportion
AMYLOID_STATUS
Amyloid negative
Amyloid positive
APOE4_binary
Carrier
Non-carrier

Supplementary Figure 8. Distribution of AD related measurements among petVAE features based clusters. Abbreviations: Aβ – Amyloid-β, ABETA - Amyloid-β cerebrospinal fluid, TAU - tau cerebrospinal fluid, PTAU - phosphorylated tau cerebrospinal fluid, PACC – preclinical Alzheimer cognitive composite, SUMMARY_SUVR – composite standardized uptake value ratio, *APOE* – Apolipoprotein E gene, Aβ-/Aβ-+/Aβ+/Aβ++ denote petVAE-based hierarchical clustering–defined groups representing increasing levels of Aβ load on PET imaging and corresponding positions along the Alzheimer's disease continuum

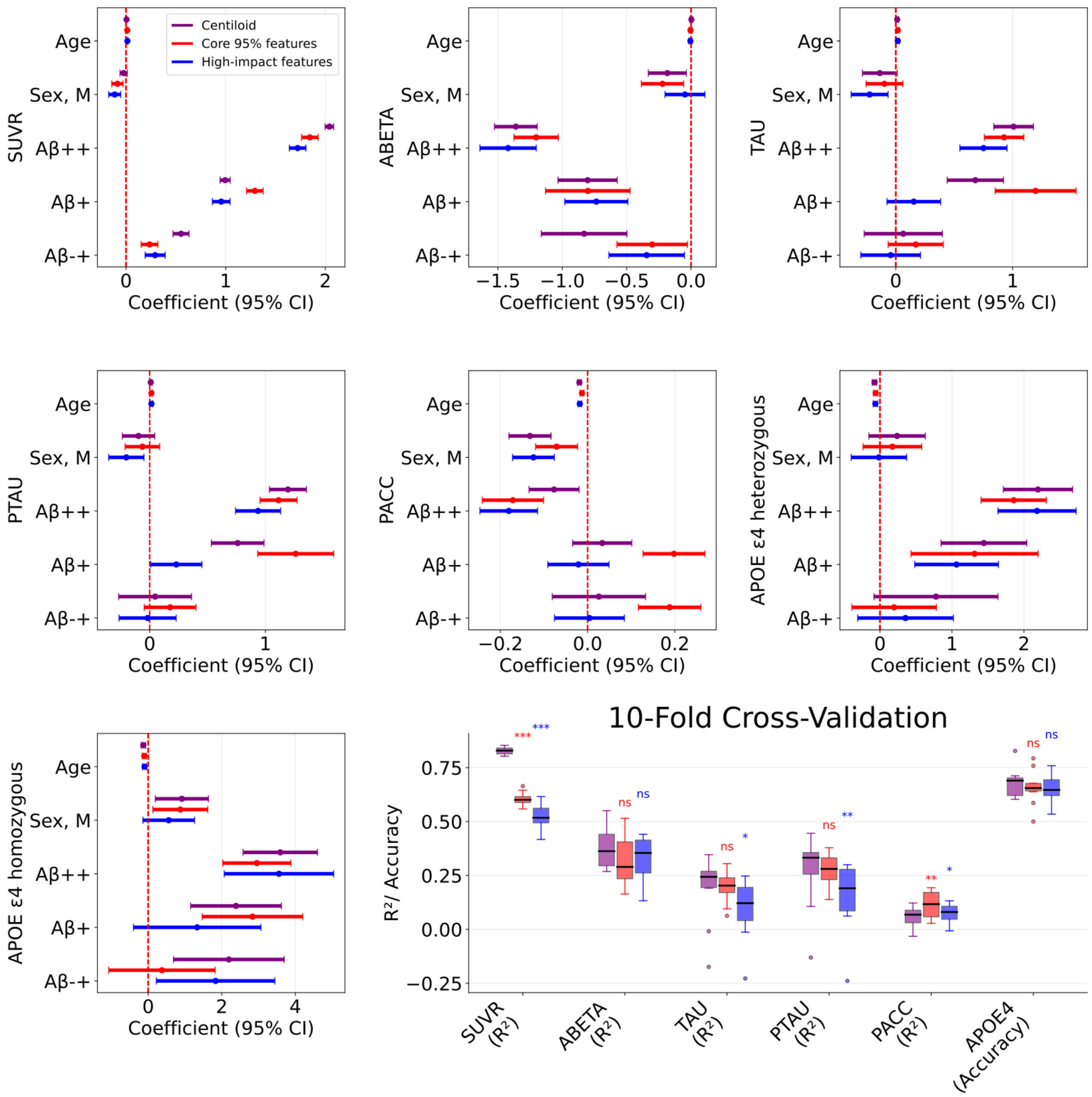

Supplementary Figure 9. Comparison of clusters by AD-related biomarkers for core 95% features, high-impact features and Centiloid scale. The Aβ- group was used as the reference group. Abbreviations: SUVR – composite standardized uptake value ratio, ABETA – Aβ cerebrospinal fluid biomarker, PACC – Preclinical Alzheimer's Cognitive Composite score, TAU – tau cerebrospinal fluid biomarker, PTAU – Phosphorylated tau cerebrospinal fluid biomarker, APOE – Apolipoprotein E gene, Aβ – Amyloid-β , CI – confidence interval,, ns - not significant, $R^2$-coefficient of determination, * - p-value< 0.05, ** - p-value< 0.01, *** - p-value< 0.001, Aβ-/Aβ-+/Aβ+/Aβ++ denote petVAE-based hierarchical clustering–defined groups representing increasing levels of Aβ load on PET imaging and corresponding positions along the Alzheimer's disease continuum.

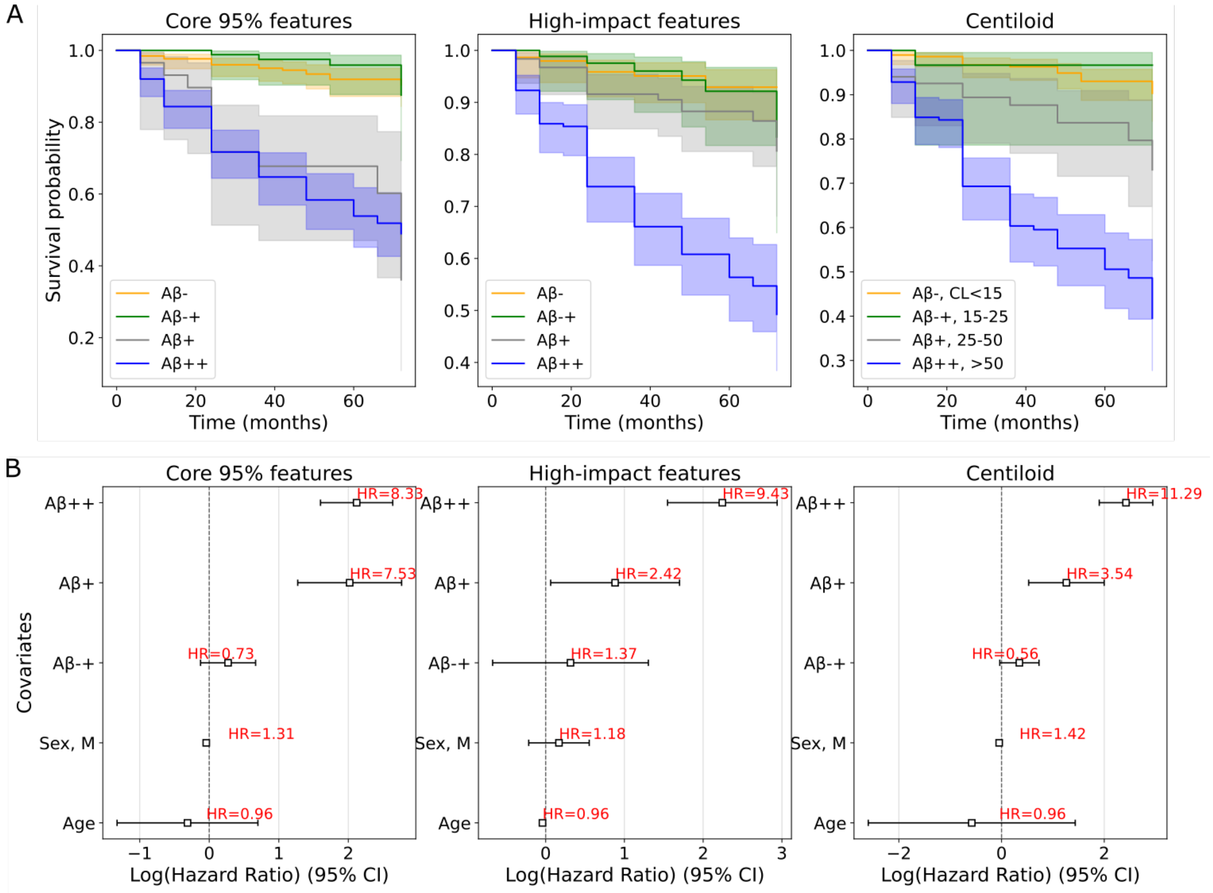

Supplementary Figure 10. Survival analysis of petVAE-based clusters for 95%-importance set, high-impact set and Centiloid scale. (A) Kaplan–Meier estimated survival curves. (B) Comparison of clusters by Cox proportional hazards models with age, sex as covariates. Aβ – Amyloid-β , HR – hazard ratio, CI – confidence interval, CL - Centiloid.

Supplementary Equation 1

$$\mathcal{L} = \mathrm{MSE}(\mathrm{x},\hat{\mathrm{x}}) + \beta\, D_{\mathrm{KL}}\big(q(z|x)\,||\,p(z)\big),$$

where L – total loss, $x$ – input, $\hat{x}$ – reconstruction, $q(z|x)$– encoder distribution, $p(z)$ – prior distribution (standard normal), and β = 0.00001 is the weighting parameter.

Supplementary Equation 2

$$\mathrm{MSE}(x,\hat{x}) = \frac{1}{N}\sum_{i=1}^{N}(x_i - \hat{x}_i)^2 \ ,$$

where $N$ is the number of pixels.

Supplementary Equation 3

$$D_{\mathrm{KL}}\big(N(\mu,\sigma^2)\ ||N(0,1)\big) = \frac{1}{2}\sum_{j=1}^{d}\big(\mu_j^2 + \sigma_j^2 - \log\sigma_j^2 - 1\big) \ ,$$

where $d$ is the dimensionality of the latent space, and $\mu_j$ and $\sigma_j^2$ are the mean and variance of the estimated posterior of the latent dimension $j$ (Kingma & Welling, 2019)

Supplementary Equation 4

$$\mathrm{SSIM}(x,y) = \frac{(2\mu_x\mu_y + C_1)(2\sigma_{xy} + C_2)}{(\mu_x^2 + \mu_y^2 + C_1)(\sigma_x^2 + \sigma_y^2 + C_2)} \ ,$$

where $\mu$ is the mean, $\sigma^2$ is the variance, $\sigma_{xy}$ is the covariance, and $C_1$, $C_2$ are small constants for stability (Wang et al., 2004). SSIM ∈ [0,1], where 1 means a perfect match with the original image and 0 means no structural similarity.

Supplementary Equation 5

$$\mathrm{PSNR}(x, \hat{x}) = 10 \cdot \log_{10}\left(\frac{L^2}{\mathrm{MSE}(x, \hat{x})}\right),$$

where L is the maximum possible pixel intensity value (*e.g.*, 1 for normalized images, 255 for 8-bit images). PSNR ∈ (−∞, +∞), where <20 dB - poor quality, 20-30 dB - moderate, > 30 dB – good quality.

Supplementary Equation 6-8

$$Concordance = \frac{(a+d)}{n}, \quad (6)$$

$$Positive\ agreement = \frac{a}{(a+b)}, \quad (7)$$

$$Negative\ agreement = \frac{d}{(c+d)}, \quad (8)$$

where a, b, c and d are the number of scans where (a) both methods assign scan to Aβ+, (b) SUVR to Aβ+, petVAE to Aβ-, (c) SUVR to Aβ-, petVAE to Aβ+, (d) both to Aβ-, and n=a+b+c+d is the total number of scans.
